# Regulation of human trophoblast gene expression by endogenous retroviruses

**DOI:** 10.1101/2022.04.26.489485

**Authors:** Jennifer M. Frost, Samuele M. Amante, Hiroaki Okae, Eleri M. Jones, Brogan Ashley, Rohan M. Lewis, Jane K. Cleal, Matthew P. Caley, Takahiro Arima, Tania Maffucci, Miguel R. Branco

## Abstract

The placenta is a fast-evolving organ with large morphological and histological differences across eutherians, but the genetic changes driving placental evolution have not been fully elucidated. Transposable elements, through their capacity to quickly generate genetic variation and affect host gene regulation, may have helped to define species-specific trophoblast gene expression programmes. Here, we assessed the contribution of transposable elements to human trophoblast gene expression as enhancers or promoters. Using epigenomic data from primary human trophoblast and trophoblast stem cell lines, we identified multiple endogenous retrovirus families with regulatory potential that lie close to genes with preferential expression in trophoblast. These largely primate-specific elements are associated with inter-species gene expression differences, and are bound by transcription factors with key roles in placental development. Using genetic editing we demonstrated that several elements act as transcriptional enhancers of important placental genes, such as *CSF1R* and *PSG5*. We also identified an LTR10A element that regulates *ENG* expression, affecting secretion of soluble ENG, with potential implications for preeclampsia. Our data show that transposons have made important contributions to human trophoblast gene regulation, and suggest that their activity may affect pregnancy outcomes.

## Introduction

The success of human pregnancy depends on the healthy development and function of the placenta. Establishment of appropriate blood flow and subsequent nutrient exchange is a carefully orchestrated balance between the requirements of the fetus and the mother, regulated by the interplay between immunological, genetic and hormonal systems at the feto-maternal interface. Following implantation, fetally derived trophoblast cells invade maternal tissues interstitially, remodelling uterine spiral arteries well into the myometrium ^1^. Aberrations to this process result in serious pregnancy complications that cause maternal and fetal morbidity and mortality, including recurrent pregnancy loss, fetal growth restriction, preterm birth and preeclampsia in the case of too little invasion, or disorders of the placenta accreta spectrum where invasion is too extensive ^2^. However, the genetic determinants of these disorders remain unclear, as genome-wide association studies have revealed very few candidates, with the notable exception of *FLT1* in preeclampsia ^3^.

Placental development and structure displays wide variation across eutherian species, even within the primate order ^4^. Notably, the deep interstitial trophoblast invasion observed in humans is unique to great apes ^5^. Placentas differ in cellular composition, histological arrangement and gross morphology, as well as many molecular aspects, all of which shape interactions between conceptus and mother, and their outcomes. This striking variation reflects the myriad selective pressures associated with the feto-maternal conflicts that fuel fast evolution of this organ ^6^. Yet, the genetic drivers of placental evolution remain to be fully elucidated.

One important yet understudied source of genetic variation is transposable elements (TEs). These abundant repetitive elements, which include endogenous retroviruses (ERVs), have made major contributions to human evolution, helping to shape both the coding and regulatory (non-coding) landscape of the genome. Akin to the variable and species-specific development and structure of the placenta, TEs are highly species-specific, making them putative drivers of placental evolution. Indeed, multiple genes with key roles in placentation have been derived from TEs ^7^, most prominently the syncytin genes, whose products mediate cell-cell fusion to generate a syncytialised trophoblast layer that directly contacts maternal blood ^8^. Additionally, the non-coding portions of TEs (e.g., the LTRs – long terminal repeats – in ERVs) have the ability to regulate gene transcription, and through this action contribute to human embryonic development ^9–11^, innate immunity ^12^, the development of cancer ^13^, and evolution of the feto-maternal interface, amongst others ^14,15^. TEs can recruit host transcription factors, often in a highly tissue-specific manner, and gain epigenetic hallmarks of gene regulatory activity (e.g., open chromatin, enrichment of H3K27ac), acting as transcriptional promoters or distal enhancer elements ^14–16^. We and others have previously shown that in mouse trophoblast stem cells, several ERV families are enriched for binding of key stemness factors (CDX2, ELF5, EOMES) and can act as major enhancers of gene expression ^17,18^. In humans, several examples of TE-encoded placenta-specific promoters have been uncovered throughout the years, such as those driving expression of *CYP19A1*, *NOS3* and *PTN* genes ^19^. A fascinating example of a human placental TE-derived enhancer has also been described, wherein a THE1B ERV regulates the expression of the corticotropin-releasing hormone, affecting gestational length when inserted into the mouse genome ^20^. More recently, the Macfarlan lab has used epigenomic data to identify a group of putative lineage-specific placental enhancers that are derived from ERVs ^21^. However, as the placenta is a heterogeneous tissue, it remains unclear whether all of these ERVs are active in trophoblast cells, and a genetic demonstration of their regulatory action is lacking.

Here, we identified ERV families that exhibit hallmarks of gene regulatory activity in human trophoblast. We show that these ERVs bind transcription factors required for placental development, and lie close to genes with preferential trophoblast expression in a species-specific manner. Using genetic editing, we show examples of ERVs that act as gene enhancers in trophoblast, including an LTR10A element within the *ENG* gene that regulates the secretion of soluble ENG protein by the syncytiotrophoblast, which is both a marker for, and contributor to, the pathogenesis of preeclampsia.

## Results

### Primate-specific ERVs exhibit regulatory characteristics in human trophoblast

To identify interspersed repetitive elements bearing hallmarks of activating regulatory potential in human trophoblast, we performed H3K27ac profiling using either ChIP-seq or CUT&Tag ^22^. We analysed our previously published data from primary human cytotrophoblast ^23^, as well as newly generated data from cytotrophoblast-like human trophoblast stem cells (hTSCs) ^24^, which can be differentiated in vitro and allow for easy genetic manipulation (Figure 1A). Using the Repeatmasker annotation (excluding simple and low complexity repeats), we determined the frequency of H3K27ac peaks per repeat family, and compared it to random controls using a permutation test (Figure 1A). This revealed 29 repeat families enriched for H3K27ac peaks in both primary cytotrophoblast and hTSCs, the vast majority of which were primate-specific ERVs (Figure 1B; Supplementary Figure 1A; Supplementary Table S1). For comparison, we performed the same analysis on published H3K27ac CUT&Tag data from human embryonic stem cells (hESCs) ^22^. Most hTSC-enriched repeats displayed little to no enrichment in hESCs (Figure 1B), despite the fact that hESCs make use of a large set of TEs for gene regulatory purposes ^10,25^. This suggests a divergence in TE-associated regulatory networks that is set up early in development, and that may help direct extraembryonic development. A more detailed analysis of H3K27ac signals across repeats belonging to each of the enriched families confirmed the asymmetry between hTSCs and hESCs (Figure 1C). HERVH-associated LTRs present an interesting case in which LTR7C elements are H3K27ac-enriched in hTSCs, whereas the related LTR7 family is H3K27ac-enriched in hESCs (Supplementary Figure 1B). Each of the LTR7 subfamilies displays a unique combination of transcription factor binding motifs ^26^, which may underlie their cell type-specific expression.

**Figure 1.**
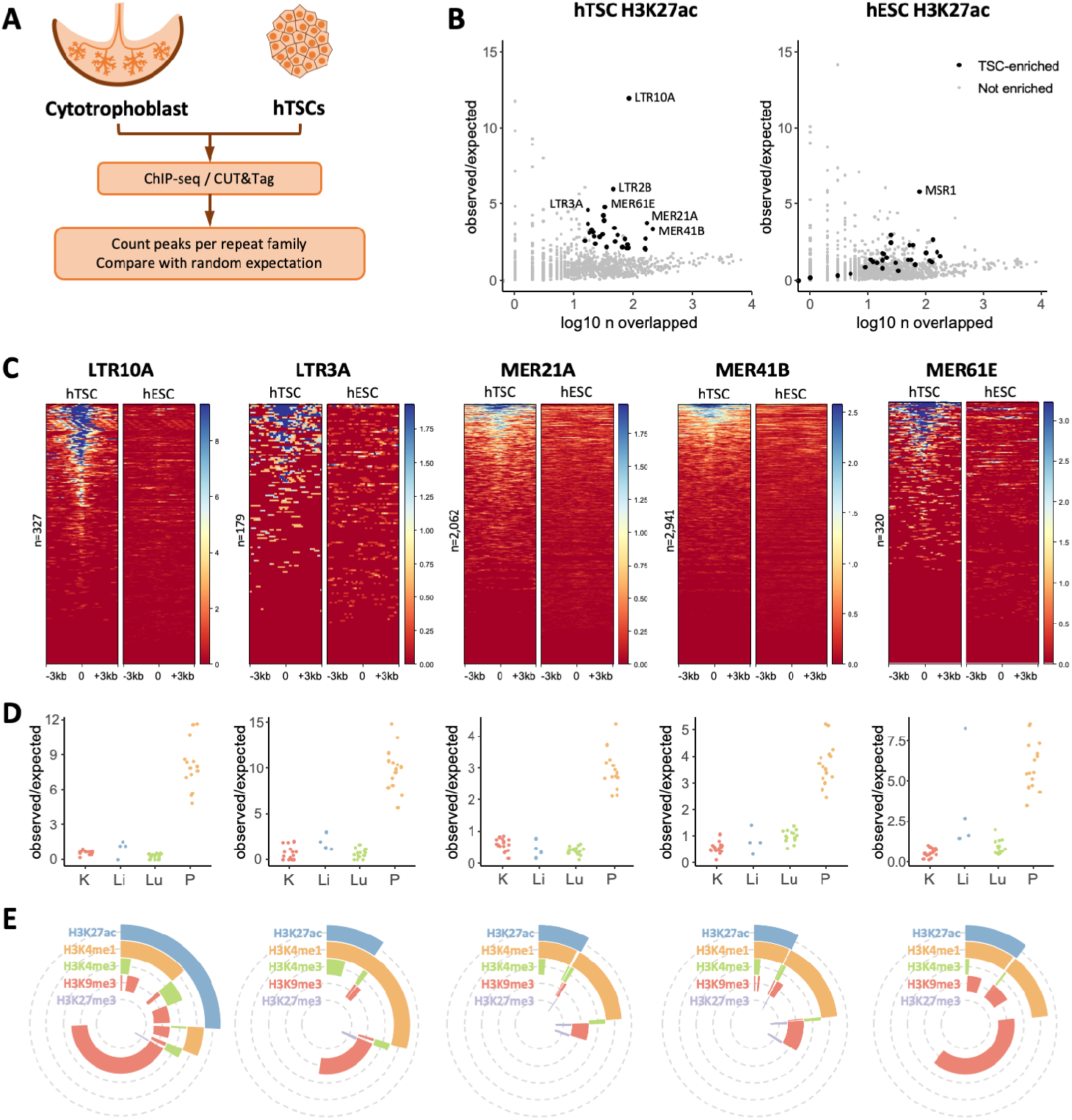
Gene regulatory signatures of ERVs in human trophoblast. A) Primary cytotrophoblast and human trophoblast stem cells were profiled for H3K27ac to identify repeat families with putative gene regulatory potential. B) Enrichment for H3K27ac peaks for each repeat family in hTSCs or hESCs. Families with significant enrichment in hTSCs are highlighted. C) H3K27ac profiles of a subset of hTSC-enriched ERV families in hTSCs and hESCs. Each line represents an element in that family. D) Enrichment for DNase hypersensitive sites in the same ERV families, in kidney (K), liver (Li), lung (Lu) and placenta (P). Each datapoint represents a different ENCODE dataset. E) Proportion of elements from the same ERV families overlapping particular combinations of histone modifications.

To further validate our findings and narrow down the list of candidate regulatory repeat families, we also analysed placental DNAse-seq data from ENCODE. We compared these data to DNAse-seq profiles from liver, lung and kidney, highlighting that most hTSC-associated families displayed tissue-specific enrichment of elements with accessible chromatin (Figure 1D; contrast with non-tissue-specific examples in Supplementary Figure 1C). Based on stringent criteria (see Methods), we decided to focus on 18 placenta-specific candidate regulatory repeat families, all of which were ERV-associated LTRs (Supplementary Table S1). This included families with known examples of elements bearing promoter activity in the placenta: LTR10A (*NOS3* gene), LTR2B (*PTN* gene), MER39 (*PRL* gene), MER39B (*ENTPD1* gene) and MER21A (*HSD17B1* and *CYP19A1* genes) ^19^.

We then characterised in more detail the H3K27ac-marked elements from each of these families by performing CUT&Tag for H3K4me1, H3K4me3, H3K9me3 and H3K27me3 in hTSCs. This revealed that a large proportion of H3K27ac-marked elements across all families was also marked by H3K4me1 (median 72%, range 44-85%), a signature of active enhancers (Figure 1E). Only a small proportion (median 3%, range 0-20%) was marked by H3K4me3, a signature of active promoters (Figure 1E). There was also a large fraction of elements marked by H3K4me1 alone (median 71% of all H3K4me1 elements, range 27-81%), which is normally associated with poised enhancers (Figure 1E), raising the possibility that this group of elements becomes active upon differentiation of hTSCs. To test this, we differentiated hTSCs into extravillous trophoblast (EVT; Supplementary Figure 1D) and performed CUT&Tag for H3K27ac. Half of the hTSC-active ERV families remained H3K27ac-enriched in EVT, whereas others displayed a clear specificity for the stem cell state (e.g., LTR10A, MER61E; Supplementary Figure 1E). In line with our hypothesis, a large proportion of ERVs that are active in EVT were found to be in a poised enhancer state in hTSCs (Supplementary Figure 1F,G). It is possible that a different set of poised ERV enhancers become active upon differentiation into syncytiotrophoblast (SynT; Supplementary Figure 1D), but the multinucleated nature of these cells seemingly interfered with our CUT&Tag attempts.

Our analyses suggest that a large number of ERVs (nearly all of which are primate-specific) may act as gene regulatory elements in human trophoblast, showing dynamic changes during differentiation.

### hTSC-active ERVs bind key placenta-associated transcription factors

Enhancers function to regulate gene expression through the binding of transcription factors, providing exquisite tissue-specific control through sequential binding of transcription factor combinations. We therefore identified transcription factor binding motifs that were enriched within each hTSC-active ERV family, and focused on a selection of transcription factors that are expressed in trophoblast (see Methods). Reassuringly, we identified previously described motifs on MER41B for STAT proteins and SRF (Supplementary Figure 2A) ^12,21^. We further uncovered a large collection of motifs for transcription factors with known roles in trophoblast development. Namely, multiple families bore motifs for key factors involved in the maintenance of the stem cell state, such as ELF5, GATA3, TFAP2C, TP63 and TEAD4 (Figure 2A; Supplementary Figure 2A). Additional transcription factors with known roles in placental development and/or physiology included JUN/FOS ^27,28^, PPARG/RXRA ^29^, and FOXO3 ^30^. Most of these motifs were also present in elements negative for H3K27ac (Supplementary Figure 2B), suggesting that there are additional genetic or epigenetic determinants of their activity, similar to what we had observed in mouse TSCs ^18^. There were nonetheless some notable exceptions where motifs were only found in active elements, such as GATA3 in LTR2B elements and SRF in MER61D elements (Supplementary Figure 2B).

**Figure 2.**
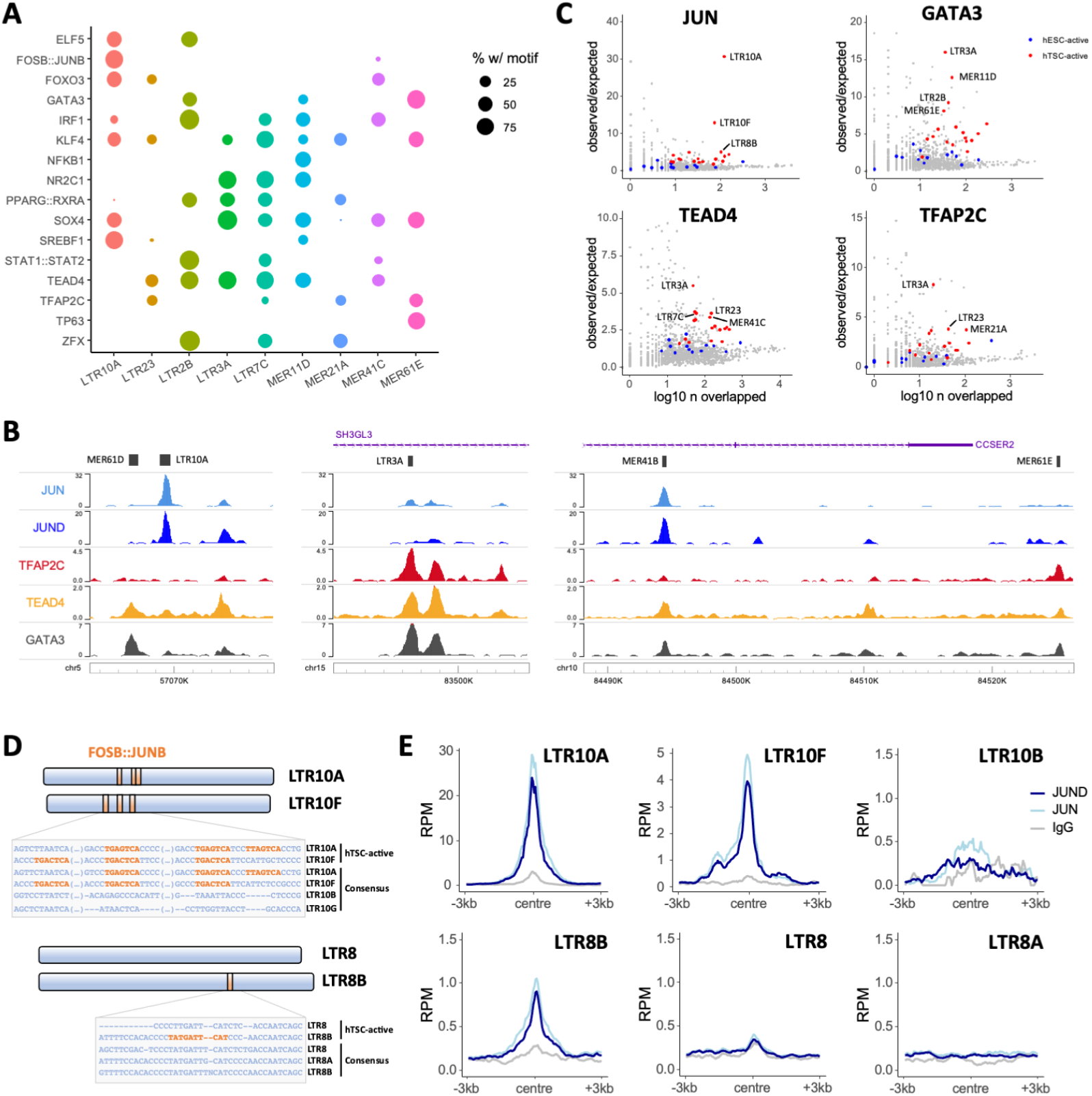
Transcription factor repertoire at hTSC-active ERVs. A) Proportion of H3K27ac-marked elements from selected ERV families bearing motifs for the transcription factors on the y axis. B) Genome browser snapshots of transcription factor CUT&Tag/RUN data, showing examples of enrichment over ERVs. C) Repeat family-wide enrichment for peaks from transcription factor CUT&Tag/RUN data on hTSCs. H3K27ac-enriched families in hTSCs or hESCs are highlighted. D) Schematic and alignment of LTR10 and LTR8 subfamilies showing the presence of FOSB∷JUNB (i.e., AP-1) motifs in the genome-wide consensus sequence and/or in a consensus of the H3K27ac-marked elements. E) Mean CUT&Tag profiles for JUN and JUND over LTR10 and LTR8 subfamilies.

To validate the binding of some of these transcription factors to hTSC-active TEs, we performed CUT&Tag or CUT&RUN ^31^ for JUN, JUND, GATA3, TEAD4 and TFAP2C (Figure 2B). We applied the same peak enrichment pipeline as used above to identify H3K27ac-enriched TE families and found that several of these families were enriched for one or more of the evaluated transcription factors (Figure 2C; Supplementary Figure 2C). In contrast, TE families active specifically in hESCs showed little to no enrichment of these factors (Figure 2C; Supplementary Figure 2C). LTR10A and LTR10F elements were strongly enriched for JUN binding, as predicted from our motif analysis. Similarly, binding to motif-bearing ERVs was confirmed for GATA3 (e.g., LTR2B, MER11D, MER61E), TEAD4 (e.g., LTR3A, LTR7C, MER41C) and TFAP2C (e.g., LTR23, MER21A). We also found instances of enriched transcription factor binding to families that seemingly do not bear the corresponding motif, such as in the case of GATA3 and TFAP2C binding at LTR3A elements. This could reflect limitations of motif-finding approaches and/or suggest that interactions between different transcription factors enable recruitment of large regulatory complexes based on a small subset of motifs – e.g., TEAD4, which binds to motifs in LTR3A elements, may recruit TFAP2C ^32^.

Given the striking enrichment for JUN and JUND binding over LTR10A/F elements (Figure 2C; Supplementary Figure 2C), we further explored the corresponding motifs in these families, as well as in LTR8B. JUN and JUND are two subunits of the AP-1 complex, which can heterodimerise with the FOS family of transcription factors. AP-1 plays important roles in cell proliferation and survival, and has been implicated in the regulation of trophoblast differentiation and invasion ^27,28^. Both LTR10A and LTR10F active elements (i.e., those marked by H3K27ac) contained three AP-1 motifs, and these were also present in the family-wide consensus sequence, whereas other LTR10-related families lacked any such motifs (Figure 2D). In strict correspondence with this, binding was observed for LTR10A/F, but not other LTR10 families (Figure 2E). It is unclear why JUN/JUND enrichment at LTR10A elements is substantially more pronounced than for LTR10F, although it could involve differences in motif arrangement, cooperative binding with other transcription factors, or chromatin environment. In the case of LTR8B, only active elements contained one AP-1 motif, suggesting divergence of a subset of elements after retroviral endogenization (Figure 2D). CUT&Tag profiles confirmed that JUN/JUND bound LTR8B only, and not other LTR8-related families (Figure 2E).

These results show that ERV families bearing regulatory potential in human trophoblast are bound by multiple transcription factors that play important gene regulatory roles in trophoblast.

### hTSC-active ERVs lie close to genes with preferential trophoblast expression

To assess the potential of hTSC-active ERVs to drive trophoblast gene expression, we first asked whether some functioned as gene promoters. Using our primary cytotrophoblast RNA-seq data ^23^, we performed de novo transcriptome assembly and extracted transcripts for which an ERV from the selected families overlapped the transcriptional start site. In line with the relative small proportion of H3K4me3-containing elements (Figure 1E), we identified few ERVs with apparent promoter activity (Supplementary Table S2). Reassuringly, this list included previously reported ERV-encoded promoters, such as those for *CYP19A1* ^33^, *PTN* ^34^, *PRL* ^35^ and *MID1* ^36^. Another notable gene was *ACKR2*, a chemokine scavenger whose expression in trophoblast is driven by a MER39 element (Supplementary Figure 3A), and deficiency of which leads to placental defects and pre/neo-natal mortality in mice ^37^. Most other transcripts associated with ERV promoters were lowly expressed in cytotrophoblast (Supplementary Table S2).

We then sought to uncover correlations between candidate ERV-derived enhancers and expression of nearby genes. We first associated each gene promoter to the nearest H3K27ac-marked TE from the selected families. Using published RNA-seq data ^24^, we then asked whether the distance to active ERVs correlated with gene expression in trophoblast cells, by comparing it to the expression in placental stroma (non-trophoblast, connective tissue containing fibroblasts and macrophages). We found a strong association between gene-ERV distance and preferential expression in both undifferentiated and differentiated trophoblast (Figure 3A), and this was also seen in data from primary cells (Supplementary Figure 3B). In contrast, genes proximal to H3K27ac-marked ERVs in hESCs displayed no preferential expression in trophoblast (Figure 3A, Supplementary Figure 3B). To further compare these two groups of genes, we analysed RNA-seq data from transdifferentiation experiments of hESCs into hTSC-like cells ^38^, which showed that genes lying within 50 kb of hTSC-active ERVs displayed higher expression upon transdifferentiation than those close to hESC-active ERVs (Figure 3B).

**Figure 3.**
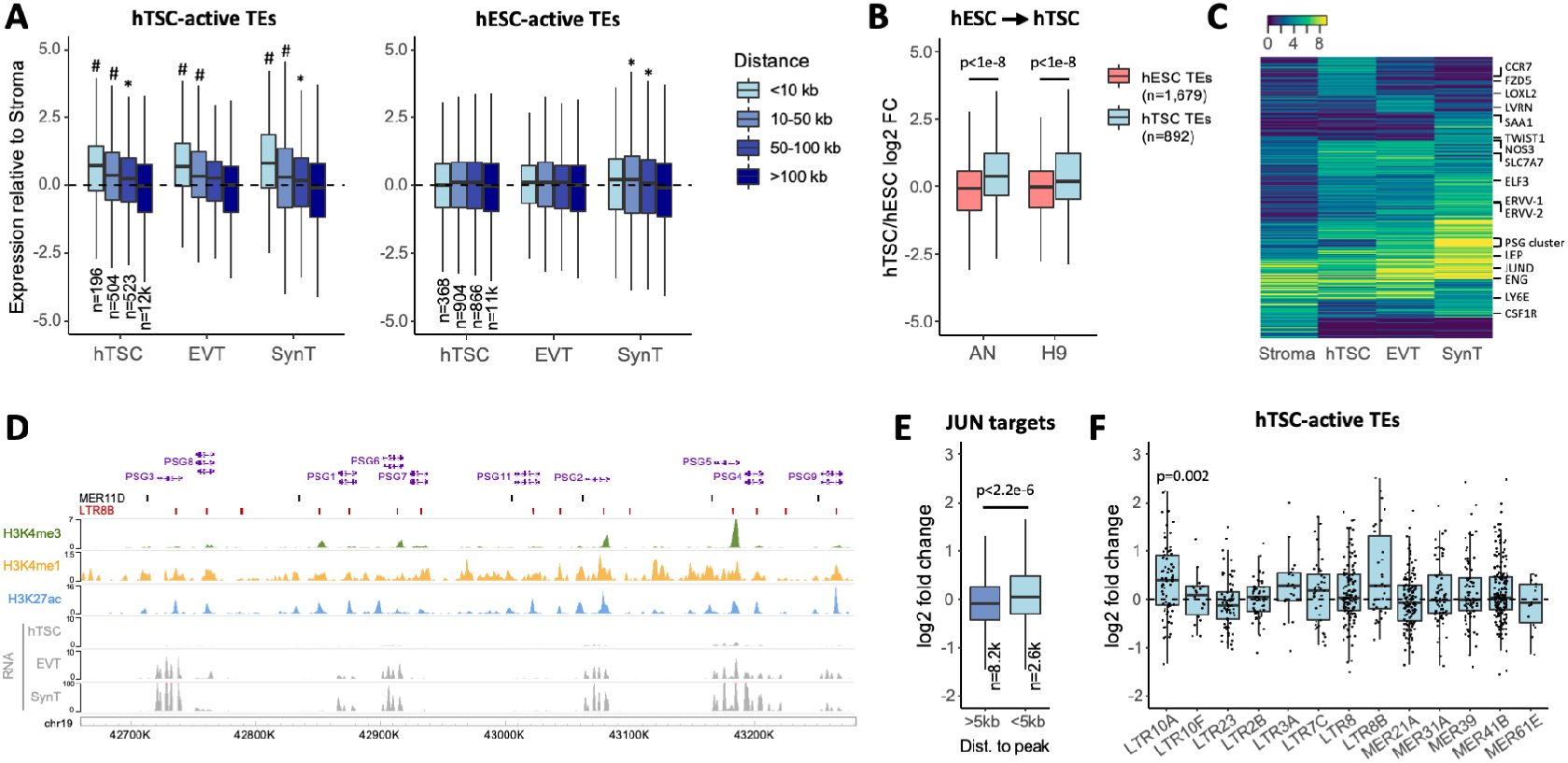
Expression of genes close to regulatory ERVs. A) Gene expression in hTSCs and hTSC-derived EVT and SynT relative to primary placental stroma, Genes are grouped based on their distance to the nearest H3K27ac-marked TE in hTSCs. The number of genes in each set is shown; * p<0.04, # p<3e-8: difference to the >100kb group based on ANOVA and Tukey post-hoc test. B) Gene expression in hESC-transdifferentiated hTSCs relative to their respective parental hESC lines (AN or H9). Expression ratios are shown for gene within 50kb or either hESC- or hTSC-active (H3K27ac-marked) TEs. P values are from Wilcoxon tests with multiple comparisons correction. C) Expression patterns of variably expressed genes across stroma and trophoblast cells. D) Genome browser snapshot of the human PSG cluster highlighting the overlap between H3K27ac peaks and MER11D or LTR8B elements. E) Fold change in gene expression after treatment of hTSCs with JNK inhibitor SP600125, with genes divided according to their distance to the nearest JUN CUT&Tag peak. The number of genes in each set is shown; p value is from a Wilcoxon test. F) Fold change in expression after SP600125 treatment for genes within 100kb of a H3K27ac-marked ERV from selected families. P value is from a Wilcoxon tests comparing each distribution to 0, with multiple comparisons correction.

Most genes proximal to hTSC-active ERVs were expressed in all three trophoblast cell types analysed, with smaller subsets displaying expression in a single trophoblast cell type (Figure 3C). A relatively large fraction showed increased expression upon differentiation into SynT. Of particular note is a cluster harbouring several genes encoding for pregnancy-specific glycoproteins (PSGs), which are highly expressed in SynT. In humans, PSGs are the most abundant conceptus-derived proteins circulating in maternal blood ^39^. Their function in pregnancy remains elusive, though they have been associated with immune responses to pregnancy, and low levels of circulating PSGs are linked to recurrent pregnancy loss, fetal growth restriction and preeclampsia ^40–42^. Within the PSG cluster, virtually every H3K27ac peak overlaps either a MER11D or LTR8B element (Figure 3D), which presumably were already present in the ancestral *PSG* gene before its duplication. These two ERV families are associated with SynT-biased gene expression, and this is largely driven by the expression of the *PSG* cluster (Supplementary Figure 3C). This is in contrast with other families that are associated with genes with increased expression across all three trophoblast cell types analysed when compared to placental stroma (Supplementary Figure 3C).

We then leveraged the information gained from our transcription factor binding analysis to interfere with putative TE regulatory target genes. Given the enrichment in JUN/JUND binding at LTR10A, LTR10F and LTR8B elements, we treated hTSCs with SP600125, an inhibitor of c-Jun N-terminal kinases (JNKs). Unexpectedly, this led to increased levels of phosphorylated c-Jun in the nucleus, and the same result was obtained with a second AP-1 inhibitor (SR11302; Supplementary Figure 4A). JNK signalling is known to have cell-type specific effects ^43^, partly due to opposing roles of JNK1 and JNK2 ^44^. JNK2 deficiency increases c-Jun phosphorylation and stability ^44^, potentially explaining our results, as JNK2 is highly expressed in hTSCs (Supplementary Figure 4B). Irrespective of the mechanism, higher levels of phosphorylated c-Jun are predicted to increase expression of AP-1 target genes. Indeed, RNA-seq of SP600125-treated hTSCs revealed that genes lying within 5 kb of a JUN binding site were on average upregulated (Figure 3E, Supplementary Figure 4D). Overall, SP600125 raised expression of a large number of genes implicated in cell migration (Supplementary Figure 4C-E), which is in line with observations that SP600125 increases trophoblast cell migration ^27^. Finally, we asked what the effects of SP600125 on predicted ERV target genes were. LTR10A target genes were upregulated upon SP600125 treatment (Figure 3F), including *NOS3* (also observed with a second AP-1 inhibitor; Supplementary Figure 4F), whose placenta-specific gene expression is driven by an LTR10A-derived promoter ^45^. Although the majority of LTR10F and LTR8B target genes were also upregulated (Figure 3F), their low number precluded robust statistical analysis. More prominent effects on LTR10A target genes were expected based on the stronger binding of JUN/JUND to this family when compared to LTR10F and LTR8B (Figure 2E). Nonetheless, expression of all *PSG* genes was increased by at least 2-fold, suggesting that JUN regulates this cluster via LTR8B elements. We also found that SP600125 led to upregulation of the internal ERV regions associated with LTR10A, but less so for LTR10F-associated ERVs (we found no LTR8B elements with a proviral arrangement), and not for other H3K27ac-enriched families (Supplementary Figure 4G), providing more direct evidence that AP-1 supports the regulatory activity of LTR10A elements.

### ERVs are associated with species-specific trophoblast gene expression

Primate evolution has involved dramatic divergence in placental phenotypes, including differences in the cellular arrangement of the feto-maternal interface and the extent of trophoblast invasion into the maternal decidua (Figure 4A) ^46^. Namely, great apes display a unique and deep form of trophoblast invasion, wherein EVT migrate both into the uterine interstitium and endothelium, remodelling spiral arteries as far as the inner third of the myometrium ^47^. The integration of TEs with regulatory capacity in trophoblast may have helped to fuel such fast placental evolution across primates. To test this, we first assessed how divergent the landscape of hTSC-active ERV families was across primate species. Using eight non-human primate genomes, we identified ERVs that were orthologous to human integrants, showing inter-species differences that are in accordance with the evolutionary age of the selected families (Figure 4B). We then took advantage of published RNA-seq data from rhesus macaque TSCs (macTSCs), which were recently derived using the same culture conditions as for hTSCs ^48^. We extracted and normalised expression values for 1-to-1 gene orthologues between human and macaque, and focused on genes within 100 kb of hTSC-active ERVs. We found that genes close to human-specific ERVs displayed on average higher expression in hTSCs than in macTSCs, when compared to genes close to conserved ERVs (Figure 4C). The majority of non-orthologous elements were from the LTR2B family (as expected from its near absence in macaque; Figure 4B) and included the previously characterised placenta-specific promoter of *PTN* ^34^, which we found to display human-specific expression when compared to macaque. Additional LTR2B-associated genes with human-specific expression included *KCNE3*, an estrogen receptor-regulated potassium channel ^49^, and *STOML2*, which regulates trophoblast proliferation and invasion ^50^.

**Figure 4.**
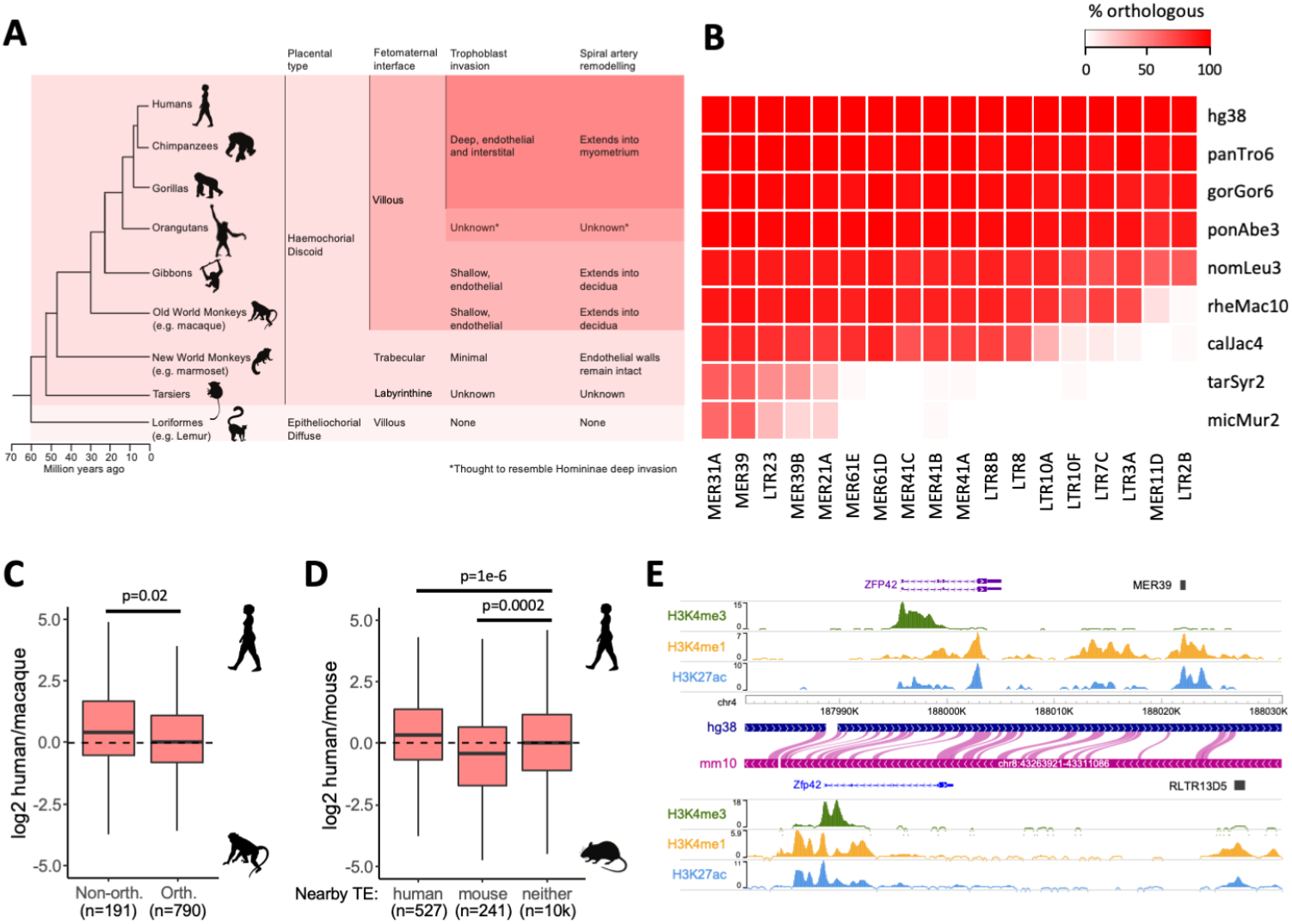
Association of ERVs to species-specific gene expression. A) Primate phylogeny highlighting cross-species differences in placental morphology and invasion^4, 47^. B) Proportion of human ERVs from each of the selected families that contains orthologous elements in the non-human primates highlighted in A. C) Expression difference between human and macaque TSCs for genes within 100kb of a H3K27ac-marked ERV from hTSC-active families, depending on whether there is an orthologous element in macaque. P value is from a Wilcoxon test. D) Expression difference between human and mouse TSCs for genes within 50kb of a H3K27ac-marked ERV in human, mouse (RLTR13D5 and RLTR13B families), or neither species. P values are from an ANOVA with Tukey post-hoc test. E) Genome browser snapshot showing a putative example of convergent evolution between mouse and human, wherein different enhancer-like ERVs lie downstream of the the *ZFP42/Zfp42* gene.

We and others have previously shown that Mus-specific ERVs also act as distal enhancers in mouse TSCs (mTSCs) ^17,18^. We therefore extended our comparative expression analysis to *Mus musculus*, asking whether human-and/or mouse-specific ERVs were associated with increased gene expression in the respective species. Indeed, genes with active ERVs nearby in human but not in mouse displayed higher expression in hTSCs, whereas those close to active mouse ERVs had higher expression in mTSCs (Figure 4D). Mouse-specific genes associated with active ERVs included a component of the FGF signalling pathway (*Fgfbp1*), which maintains the stem cell state in mouse but not in human trophoblast. Conversely, one human-specific ERV (MER41B) was associated with expression of a Wnt signalling receptor (*FZD5*), a pathway that is important for hTSC derivation ^24^. Other ERV-associated mouse-specific genes included *Duox*, *Duox2* and *Nr0b1*, which are mTSC markers ^51^, and in human *MMP14*, which is important for trophoblast invasion ^52^. We also considered potential cases of convergent evolution, whereby the same gene may be regulated by different ERVs in human and mouse. Out of the 12 genes that were close to active ERVs in both mouse and human, 10 were expressed in mTSCs and hTSCs, including *Zfp42*/*ZFP42* (Figure 4E; Supplementary Table S3). Despite being a well-known marker of ESCs, *Zfp42* is also expressed in mouse trophoblast, especially in early embryos ^53,54^, where it regulates the expression of some imprinted genes ^53^. This raises the possibility that different ERVs have convergently been co-opted to maintain the expression of *Zfp42*/*ZFP42* in mouse and human trophoblast, to support its essential roles therein. These analyses suggest that some of the putative regulatory ERVs that we identified help to drive species-specific expression of genes that are important for trophoblast development and function.

### ERVs regulate the expression of genes involved in human trophoblast function

Our data demonstrate that many primate-specific ERVs exhibit epigenetic characteristics of regulatory elements in human trophoblast. However, previous observations have shown that these markers are not predictive of gene regulatory activity ^18^. To test whether ERVs can act as enhancers in vivo, we utilised CRISPR to genetically excise a subset of candidate regions, and then measured nearby gene expression. Because the efficiency of growing clonal hTSCs from single cells was extremely low after CRISPR, we employed a population-wide lentiviral approach that we previously used ^55^, achieving an average of 49% deletion efficiency across different targets and experiments (Supplementary Table S4).

We first excised an enhancer-like MER41B element that is conserved from New World monkeys to humans, and is located in the first intron of the *ADAM9* gene (Figure 5A), which encodes for a metalloproteinase. Genetic variants of *ADAM9* are implicated in preeclampsia ^56^, and its known substrates play roles in inflammation, angiogenesis, cellular migration and proliferation ^57^. Two independent MER41B excisions were derived (603 and 751bp), resulting in a 1.7-2 fold decrease in *ADAM9* expression in hTSCs compared to no-sgRNA controls (Figure 5A). The MER41B LTR is also 8 kb upstream of *TM2D2* and 16 kb upstream of *HTRA4*, a placenta-specific serine peptidase that is upregulated in early-onset preeclampsia ^58,59^. Expression of these genes was low and remained largely unchanged following MER41B excision (Supplementary Figure 5A).

**Figure 5.**
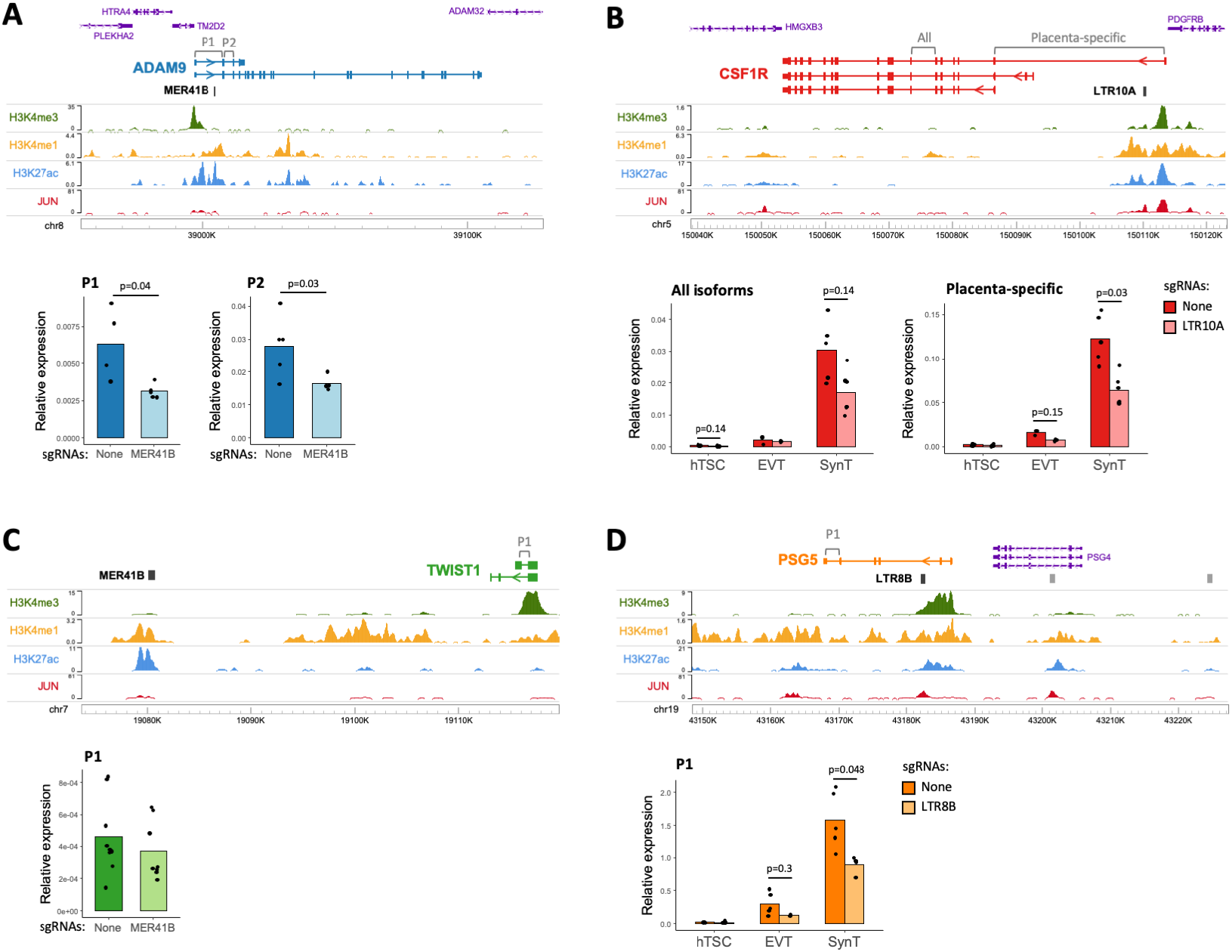
Genetic excision of hTSC-active ERVs. A) Genome browser snapshot of the ADAM9 locus containing an enhancer-like MER41B element, and ADAM9 expression (using primer pairs highlighted on the gene annotation) in hTSC populations treated with lentiviral *CRISPR* constructs carrying either no sgRNAs or sgRNAs that excise the MER41B element (n=4-5 from two sgRNA sets). B) As in A, but for CSF1R and its intronic LTR10A elements (n=3-6 from one sgRNA set). CSF1R expression was measured in hTSCs and hTSC-derived EVT and SynT. C) As in A, but for *TWIST1* and a downstream MER41B element (n=8-9 from three infections using one sgRNA set). D) As in A, but for *PSG5* and its intronic LTR8B elements (n=3-6 from two sgRNA sets). *PSG5* expression was measured in hTSCs and hTSC-derived EVT and SynT. P values throughout are from Wilcoxon tests with multiple comparisons correction.

The *CSF1R* gene has a placenta-specific promoter ^60^, downstream of which lies an LTR10A that features enhancer-like chromatin features and JUN binding in hTSC (Figure 5B), and is conserved from Old World monkeys to humans. Both *CSF1* and *CSF1R* expression increase in the placenta during pregnancy ^61^. CSF1 signalling via CSF1R promotes the growth, proliferation and migration of trophoblasts in humans and mice ^62–64^, and high CSF1 levels are correlated with preeclampsia development ^65^. Expression of both *CSF1R* transcript variants was low in undifferentiated hTSCs, but increased following differentiation to EVT, and was highest in SynT-differentiated hTSCs, particularly for the placenta-specific *CSF1R* variant (Figure 5B). *CSF1R* expression also increased upon SP600125 treatment, suggesting it is regulated by AP-1 (Supplementary Figure 4F). A 693 bp excision of the LTR10A was derived in hTSCs, with *CSF1R* expression of the placenta-specific variant being reduced by around 2-fold in both EVT and SynT cell pools (and to a lesser extend in hTSCs), compared to no-sgRNA controls (Figure 5B). These differences were not caused by an impairment in trophoblast differentiation efficiency, as judged by the expression of key marker genes (Supplementary Figure 5C). The LTR10A element therefore acts as an important enhancer of *CSF1R* in human trophoblast.

We excised a second MER41B element with enhancer-like chromatin conformation (Figure 5C), deriving a 1,469 bp deletion in hTSCs. The *TWIST1* (35 kb downstream), *FERD3L* (65 kb downstream) and *HDAC9* (79 kb upstream) genes lie in the vicinity of this LTR, but only *TWIST1* is expressed in hTSCs. TWIST1 regulates the syncytialisation of trophoblast, perhaps via GCM1 ^66,67^ and promotes epithelial to mesenchymal transition – a key process in EVT differentiation ^68^. We measured expression of *TWIST1* in MER41B excision hTSC pools, and found its expression to be unchanged (Figure 5C). Since TWIST1 is important for trophoblast differentiation, we also differentiated hTSCs to EVT and SynT, resulting in an 25-41 fold increase in *TWIST1* expression, but there was no difference in *TWIST1* expression in excision versus no-sgRNA differentiated cells (Supplementary Figure 5B). This particular MER41B element is therefore either a redundant enhancer or does not regulate *TWIST1*, highlighting the importance of these genetic experiments.

As previously mentioned, the *PSG* cluster on chromosome 19 includes MER11D and LTR8B elements at each tandemly repeated gene locus, all featuring enhancer-like chromatin features in hTSCs (Figure 3D). In order to test LTR functionality, we excised an LTR8B element, conserved only in apes, within the second intron of one of the most highly expressed PSG in humans, *PSG5* (Figure 5D) ^69^, deriving two independent excisions (774 and 901 bp). *PSG5* expression was low in undifferentiated hTSCs and remained unchanged in LTR8B excision pools compared to no sgRNA controls (Figure 5D). However, following differentiation to EVT and SynT, *PSG5* expression was increased by 15 and 81 fold respectively, and was reduced in LTR8B excision pools compared to no-sgRNA controls by 2.3 fold in EVT and 1.8 fold in SynT (Figure 5D), whilst differentiation efficiency was unaffected by the excision (Supplementary Figure 5D). These data indicate that the *PSG5* LTR8B element acts an enhancer for *PSG5* expression in human EVT and SynT. The fact that the enhancer activity of this LTR8B-*PSG5* element and the LTR10A-*CSF1R* element are most strongly expressed after differentiation supports the notion that some hTSC-active ERVs also play roles (and some may be poised to do so) in differentiated trophoblast, as suggested by our epigenomic and transcriptomic analyses above.

### An LTR10A-derived enhancer promotes ENG expression, affecting sENG secretion

We were particularly interested in an enhancer-like (and JUN-bound) LTR10A element within the first intron of the Endoglin gene (*ENG*/*CD105*) (Figure 6A). ENG is a transforming growth factor-beta (TGF-ß) 1 and 3 co-receptor, with both membrane-bound and soluble cleavage variants, highly expressed in the endothelium and SynT, and involved in the pathogenesis of preeclampsia ^70^. Indeed, the serum levels of soluble ENG (sENG) are strongly correlated with the severity of preeclampsia ^71^. Membrane-bound ENG is also expressed in villous cell columns in the first trimester of pregnancy, regulating trophoblast differentiation from proliferative cells to the migratory, invasive EVT that colonise the uterus and remodel maternal spiral arteries ^72,73^.

**Figure 6.**
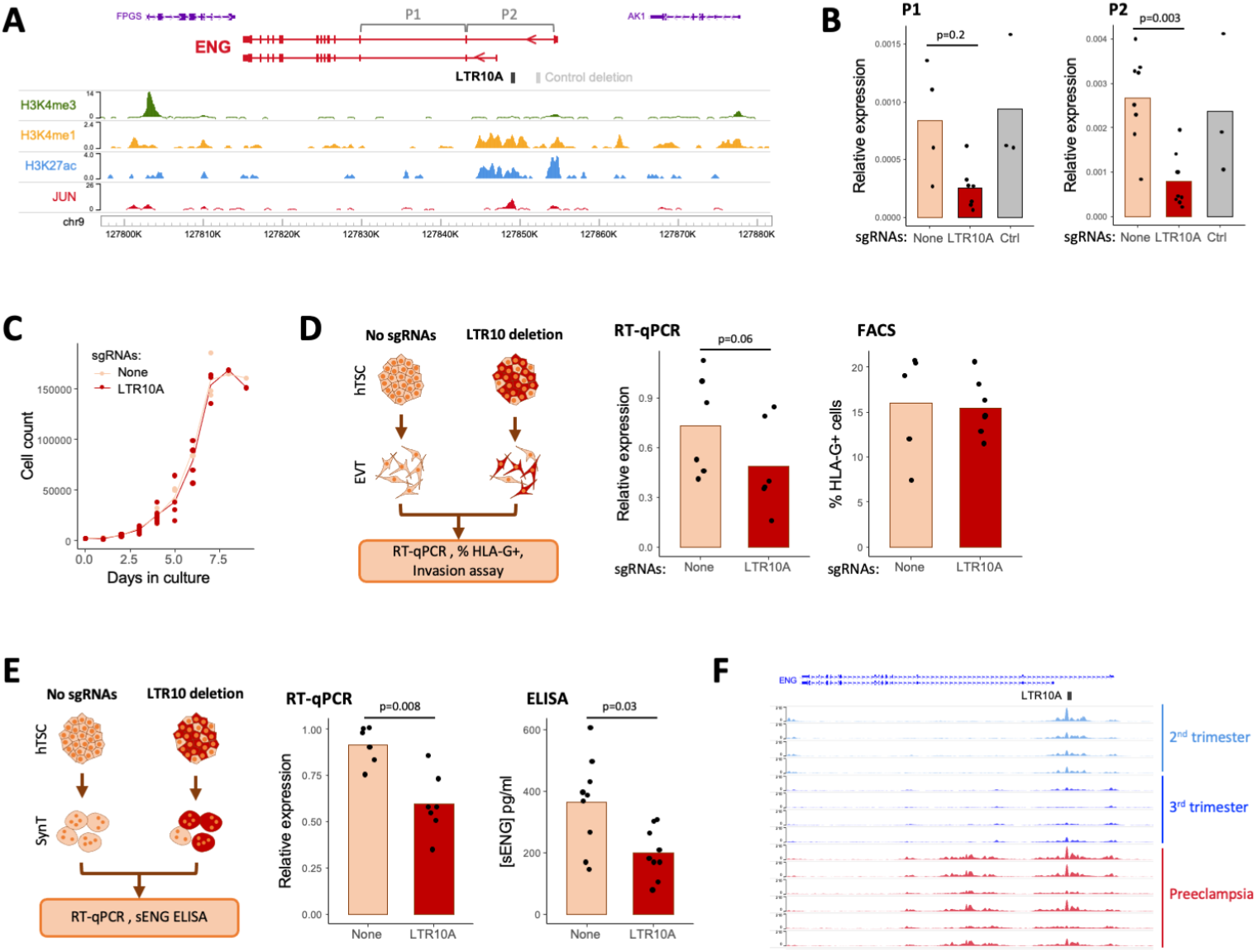
Impact of an LTR10A element within the *ENG* gene. A) Genome browser snapshot of the *ENG* locus containing an enhancer-like LTR10A element in its first intron. B) *ENG* expression (using primer pairs highlighted in A) in hTSC populations treated with lentiviral CRISPR constructs carrying no sgRNAs, sgRNAs that excise the LTR10A element, or sgRNAs that excise a control region highlighted in A (n=3-7 from three sgRNA sets). C) Growth curves for hTSC populations carrying no sgRNAs or LTRIOA-excising sgRNAS. D) The same hTSC populations were differentiated to EVT and assays performed for ENG RNA levels and HI_A-G cell surface expression. E) The same hTSC populations were differentiated to SynT and assays performed for *ENG* RNA levels and secretion of soluble ENG protein. F) Genome browser snapshot of the *ENG* locus with cytotrophoblast H3K27ac ChIP-seq data from second or third trimester uncomplicated pregnancies, or third trimester severe preeclampsia placentas. P values throughout are from Wilcoxon tests with multiple comparisons correction where relevant.

We derived three independent excisions of the *ENG* LTR10A in hTSCs (508, 614 and 874 bp), which resulted in a striking decrease in *ENG* expression compared to no-sgRNA controls (Figure 6B). Expression of neighbouring genes (*AK1* and *FPGS*) did not change (Supplementary Figure 6A). To confirm that loss of *ENG* expression was strictly associated with deletion of the LTR10A element, and not transcriptional interference by the CRISPR-Cas9 machinery, we also deleted a control region upstream of the putative LTR10A enhancer that was devoid of active histone modification marks in hTSCs (Figure 6A). Reassuringly, deletion of this region (1401 bp) in hTSCs had no effect on *ENG* expression (Figure 6B), confirming that the LTR10A element is a bona fide enhancer. In line with the role of AP-1 in regulating LTR10A regulatory activity, increased c-Jun phosphorylation led to *ENG* upregulation (Supplementary Figure 4F). To assess the phenotypic impact of LTR10A deletion, we first measured cell proliferation in hTSCs, finding no difference when compared to no-sgRNA controls (Figure 6C). We then asked whether LTR10A deletion affected trophoblast differentiation and/or phenotypes within differentiated cell types.

Differentiation to EVT was not affected by LTR10A deletion, as measured by the percentage of HLA-G-positive cells (Figure 6D; Supplementary Figure 6B). LTR10A-deleted EVT were also morphologically similar to control EVT and retained the capacity to invade Matrigel in a transwell assay (Supplementary Figure 6C), although technical variation precluded a more quantitative assessment of the extent of invasion. Notably, ENG expression remained lower (albeit variable) in LTR10A-deleted EVT, when compared to no-sgRNA controls (Figure 6D). Differentiation into SynT was also unaffected by deletion of the LTR10A element (Supplementary Figure 6D). Despite a large rise in *ENG* levels upon SynT differentiation, LTR10A-deleted cells expressed less *ENG* than no-sgRNA controls (Figure 6E). We therefore tested whether sENG protein levels were impacted by the LTR10A enhancer by performing ELISA assays in the media of day 6 differentiated SynT cultures. We detected sENG in the media at concentrations varying from ~100 pg/ml to 600 pg/ml, depending on the experiment, and observed a significant decrease in sENG protein levels in *ENG*-LTR10A excised SynT cultures when compared to no-sgRNA control (Figure 6E).

These experiments raise the possibility that deregulation of the *ENG*-LTR10A element may be associated with elevated levels of *ENG* in preeclampsia. We therefore leveraged recently published H3K27ac ChIP-seq data from cytotrophoblast isolated from preeclamptic or uncomplicated pregnancies ^74^. In uncomplicated pregnancies, there was clear H3K27ac enrichment over the LTR10A-*ENG* element in second trimester placentas, but decreased in third trimester placentas (Figure 6F). However, in third trimester placentas from severe preeclampsia, H3K27ac levels remained high over the LTR10A-*ENG* enhancer (Figure 6F), suggesting that this ERV may help to maintain ENG expression unduly high in preeclampsia.

These data show that the LTR10A-*ENG* element acts as an enhancer in human trophoblast, regulating *ENG* expression, sENG protein production, with potential implications for preeclampsia.

## Discussion

We have identified multiple ERV families that are enriched for elements bearing a chromatin signature of *cis* regulatory elements in human trophoblast, most resembling enhancers. Our stringent criteria ensured that the regulatory profiles of these families reflect what is observed in vivo and are not driven by non-trophoblast cell types present in the placenta. Indeed, we find that the activity of these families is largely trophoblast-specific, with several being active and playing important roles in both undifferentiated and differentiated trophoblast cell types. Our CRISPR genetic editing experiments demonstrated that at least a subset of the ERVs we identified by chromatin status act as bona fide enhancers of genes with important roles in placentation. Notably, the fact that we were limited to performing CRISPR on a population scale implies that the effects we observed are actually an underestimation of the true importance of those ERVs to gene expression.

We have already highlighted how many of the identified ERV families include elements that act as placenta-specific promoters ^19^. We also compared our selected families with those identified by the Macfarlan lab as being enriched for lineage-specific placental enhancers ^21^. Reassuringly, we find multiple ERV families in both studies, including MER21A, MER41B, LTR8 and MER39 ^21^. On the other hand, the Sun et al. study did not list other ERV families identified here, most prominently LTR10A, which has a strong regulatory signature (Figure 1B), and copies of which we demonstrated act as enhancers of important placental genes (Figure 5B and 6). Conversely, we find no evidence of regulatory activity from the MaLR group of ERV families identified by Sun et al. One possible reason for these discrepancies is that Sun et al. used whole placental explants, leading to a mixed epigenomic profile from multiple cell types.

We also noted that a number of our trophoblast-active ERV families were recently identified as active in several cancers, including LTR10A, LTR10F, LTR2B and MER11D ^75^. Multiple parallels have previously been drawn between trophoblast and cancer cells, including an epigenetic landscape that is strikingly different from other differentiated cell types ^76^. The combination of an arguably permissive chromatin conformation and shared signalling pathways ^77^ may make the co-option of ERVs for both placental development/function and cancer a frequent occurrence. For example, the Hippo signalling pathway, acting through YAP and TEAD4, plays key roles in both tumourigenesis and placental development ^78^. The activation of LTR10A elements in both contexts is also seemingly driven by a shared signalling pathway, in this case MAPK/AP-1 ^75^.

Several specific elements identified here suggested a potential important role for ERVs in placental evolution. This included the multiple LTR8B and MER11D copies associated with the *PSG* gene cluster (Figure 3D). The *PSG* cluster is semi-conserved in primates, with 6 to 24 genes found in Old World monkeys, 1 to 7 genes in New World monkeys and none in more distantly related primates such as lemurs, suggesting that the presence of *PSG* may correlate with haemochorial placentae, since lemurs have an epitheliochorial placenta ^79^. Notably, *PSG* clusters are also present in mice, which also have a haemochorial placenta and this region was expanded independently in mice and primates, suggesting convergent evolution ^80^. Our results suggest that the integration of LTR8B elements ahead of *PSG* cluster expansion in humans was an important step that contributed to high trophoblast expression of these genes. MER11D elements may play a similar role, and together with LTR8B be responsible for much of the transcriptional regulation of this important locus. Similarly, MER61D/E retrotransposition may have played a key role in setting the TP63 binding landscape, which in human trophoblast supports cell proliferation and prevents cellular differentiation in trophoblast ^81^. TP63 belongs to the same family of transcription factors as TP53, sharing many of its binding sites ^82^. It was previously shown that MER61 elements expanded the TP53 binding network in primates between 56 and 81 million years ago, and a number of copies have been exapted to mediate cellular stress responses in lymphoblastoid cells ^83^.

Other examples suggest a role of ERVs in coordinating the expression of genes from the same pathway. MER21A elements were previously shown to act as promoters of the steroidogenesis pathway genes *CYP19A1* and *HSD17B1*, implicating these LTRs in the regulation of steroidogenesis in human trophoblast. We also noted that both *NOS3* and *ENG* bear LTR10A elements as major transcriptional regulators (as promoter and enhancer, respectively). This is interesting because the contribution of sENG to vascular pathology in preeclampsia is partially due to effects on NOS3; in concert with FLT1, ENG reduces placental angiogenesis and vasodilation of maternal spiral arteries, and increases vessel permeability ^70,84^.

Our results implicate regulatory ERVs in pregnancy complications, prompting the need for further investigation of their functional impact on pregnancy outcomes. These ERVs may bear genetic variants that are difficult to investigate due to their repetitive nature, and that may affect their regulatory activity in the placenta. Notably, structural variants in LTR10A/F elements (in the form of variable number tandem repeats), some potentially contributing to cancer, were recently described ^75^. Research into the effects of such variants in the placenta will benefit from more complex models that can assess the impact of regulatory ERVs on cell-cell interactions, such as those between the conceptus and the mother, that make pregnancy so unique. Such experiments will be greatly supported by the recent development of placental and endometrial organoids, as well as platforms that support the study of trophoblast invasion ^85,86^.

## Materials and Methods

### Tissue culture

hTSCs were cultured according to ^24^, with modifications outlined in ^87^. Briefly, ~1 × 10^5^ hTSC were seeded onto 6 well plates coated with either 5–10 μg/mL collagen IV (10376931, Fisher Scientific) or 0.5 μg/mL iMatrix 511 (NP892-011, Generon). Basal TS medium comprised DMEM/F12 (Gibco) supplemented with 1% KnockOut Serum Replacement (KSR; Life Technologies Ltd Invitrogen Division), 0.5% Penicillin-Streptomycin (Pen-Strep, Gibco), 0.15% BSA (A9205, Sigma-Aldrich), 1% ITS-X supplement (Fisher Scientific) and 200 μM l-ascorbic acid (Sigma-Aldrich). Complete media contained 2.5μm Rho-associated protein kinase (ROCK) inhibitor Y27632, TGF-B inhibitors A83-01 (5uM), CHIR99021 (2uM) (all StemMACS) and epidermal growth factor (50ng/ml, E9644, Sigma-Aldrich), with 0.8mM histone deacetylase inhibitor Valproic Acid (PHR1061, Sigma-Aldrich). hTSC were passaged using TrypLE Express (Fisher Scientific) and split 1:3 to 1:5 every 2-3 days. All cells were cultured at 37°C in 5% CO_2_ and 20% O_2_. For JNK inhibition experiments, hTSCs were seeded onto 12-well or 6-well plates and treated with either: 10 μM SP600125 (Cambridge Bioscience) for 2-4 days, or 20 μM SR11302 (Cambridge Bioscience) for 1-2 days. Control wells were treated with DMSO. Changes in the expression of diagnostic genes were consistent across different cell collection time points.

### Differentiation to EVT and SynT

Differentiation was performed as outlined in ^24^. For EVT, hTSC were seeded in a 6-well plate coated with 1 μg/ml Col IV at a density of 0.75 × 10^5^ cells per well and cultured in 2 mL of EVT medium: DMEM/F12 supplemented with 0.1 mM 2-mercaptoethanol, 0.5% Pen-Strep, 0.3% BSA, 1% ITS-X, 100 ng/ml Human Neuregulin-1 (NRG1; Cell Signalling), 7.5 μM A83-01, 2.5 μM Y27632, and 4% KSR. Cells were suspended in the medium, and Matrigel (LDEV-free, 354234, Scientific Laboratory Supplies) added to a final concentration of 2%. On day 3, medium was replaced with EVT medium without NRG1, and Matrigel added to a final concentration of 0.5%. On day 6, medium was replaced with EVT medium without NRG1 and KSR, and Matrigel added to a final concentration of 0.5%, with cells grown for a further two days. Differentiation efficiency was measured using Fluorescence Activated Cell Sorting (FACS) following staining with an APC conjugated antibody to HLAG (APC anti-human HLA-G Antibody Clone: 87G, BioLegend UK), in unmodified hTSC this was consistently between 40-70%, and in lentivirus-infected CRISPR excision lines and controls, between 10-30 %. SynT (3D) differentiation was carried out by seeding 2.5 × 10^5^ hTSC in uncoated 6 well plates, cultured in 2 mL of SynT medium: DMEM/F12 supplemented with 0.1 mM 2-mercaptoethanol, 0.5% Pen-Strep, 0.3% BSA, 1% ITS-X, 2.5 μM Y27632, 50 ng/ml EGF, 2 μM Forskolin (Cambridge Bioscience), and 4% KSR. 2 ml fresh ST(3D) medium was added on day 3. Cells were passed through a 40 μm mesh strainer and cells remaining on the strainer were collected and analysed. Differentiation was assessed using RT-qPCR and immunostaining (Primers listed in Supplementary Table S5)

### FACS

Cells were single-cell dissociated using TrypLE Express and washed in FACS buffer (PBS supplemented with 4% KSR). The cells were then resuspended in 500 μL fresh FACS buffer, passed through a 70 μm cell strainer, and GFP+ cells collected and replated (lentiviral infections). To sort EVT differentiated cells, 400 μL of the cell suspension was incubated with APC-HLAG for 15 minutes in the fridge, 100 μL was used as an unstained control. Following antibody incubation, the cells were washed twice with FACS buffer, resuspended in fresh FACS buffer, and passed through a 70 μm cell strainer. Flow cytometry was performed using a FACS Aria II, and the data analyzed using the FlowJo software.

### Immunostaining

5000 hTSC per well were grown on collagen IV coated glass coverslips in a 24 well plate, and the differentiation protocol to EVT followed. On day 8 of EVT differentiation, media was removed and cells washed with PBS x3, before fixing in 4% paraformaldehyde for 10 minutes before staining. Differentiated SynT cells were centrifuged briefly (300g, 1 minute), resuspended gently in 500ul PBS + 4% KSR, and added dropwise to poly-L-lysine coated glass coverslips. Once SynT cells had gathered on the cover slip, the PBS+KSR was removed and cells were fixed in 4% paraformaldehyde for 10 minutes. Both fixed EVT and SynT cells were permeabilised with 0.1% Triton-X in blocking buffer (1% BSA (w/v) 2% FCS (v/v)) for 5 minutes, followed by 1 hour incubation in blocking buffer. Incubation with the primary antibody (SDC-1 (1:200, CD138 Mouse anti-Human, PE, Clone: MI15, Fisher Scientific, UK) for SynT and HLAG (1:100, APC anti-human HLA-G Antibody Clone: 87G, BioLegend UK) for EVT) diluted in blocking buffer was performed at 4°C overnight. The secondary Alexa Fluor 488-green antibody was added at a 1:250 dilution for 1 hour at room temperature. DAPI (300nM) was used as a nuclear stain. Images were collected using a Leica DM4000 epifluorescence microscope.

### Western Blot

Cells were washed with ice-cold PBS, lysed with ice-cold ‘Triton lysis buffer’ (50 mM Tris, pH 7.5, 150 mM NaCl, 1% Triton X-100, supplemented with protease inhibitor cocktail, PMSF and sodium orthovanadate) and pelleted. Supernatants were labelled as ‘Triton-soluble fractions’, which are primarily cytoplasmic. Pellets were washed with ice-cold PBS and lysed with 2% SDS, followed by sonication (these were labelled as ‘Triton-insoluble fractions’, which are primarily nuclear). Protein concentrations were assessed using Pierce™ BCA Protein Assay Kit (Thermo Fisher Scientific). Samples were separated by SDS-PAGE and transferred into nitrocellulose membranes. Membranes were incubated with 5% skimmed milk in PBS-T for 30 min at room temperature, followed by overnight incubation with the following antibodies at + 4 °C: anti pSer63 c-Jun (Cell Signalling, #2361, 1:1,000), anti c-Jun (Cell Signalling, #9165, 1:1,000). After washing with PBS-T, membranes were incubated with peroxidase conjugated anti-rabbit IgG (Sigma Aldrich, A6154, 1:10,000) for 1 h at room temperature, washed with PBS-T and exposed to ECL reagent (Sigma Aldrich, cat no: WBKLS0500). Signals were visualized using X-ray film and a film processor (pSer63 c-Jun) or using Chemidoc™ MP Imaging System (Bio-Rad). Membranes were re-incubated with anti α-Tubulin (Sigma Aldrich, T9026, 1:20,000) or anti H3 (Abcam, ab1791, 1:10,000) for 1h at room temperature, washed and then incubated for 1h with peroxidase conjugated anti-mouse IgG (Sigma Aldrich, A0168, 1:10,000) or anti-rabbit IgG, respectively, followed by visualisation using Chemidoc™.

### ChIP-seq

hTSC were dissociated using TrypLE express, pelleted and washed twice with PBS. Cell pellets were fixed with 1% formaldehyde for 12 minutes in PBS, followed by quenching with glycine (final concentration 0.125 M) and washing. Chromatin was sonicated using a Bioruptor Pico (Diagenode) to an average size of 200–700 bp. Immunoprecipitation was performed using 10 μg of chromatin and 2.5 μg of human antibody (H3K4me3 [RRID:AB_2616052, Diagenode C15410003], H3K4me1 [RRID:AB_306847, Abcam ab8895], and H3K27ac [RRID:AB_2637079, Diagenode C15410196]). Final DNA purification was performed using the GeneJET PCR Purification Kit (Thermo Scientific, K0701) and eluted in 80 μL of elution buffer. ChIP-seq libraries were prepared from 1 to 5 ng eluted DNA using NEBNext Ultra II DNA library Prep Kit (New England Biolabs) with 12 cycles of library amplification.

### CUT&Tag (Cleavage Under Targets and Tagmentation)

CUT&Tag was carried out as in ^22^. 100,000 hTS or 50,000 EvT cells per antibody were harvested fresh using TrypLE-express and centrifuged for 3 minutes at 600×g at room temperature (RT). Cells were washed twice in 1.5 mL Wash buffer (20 mM HEPES pH 7.5; 150 mM NaCl; 0.5 mM Spermidine; 1x Protease inhibitor cocktail; Roche 11836170001 (PIC)) by gentle pipetting. BioMagPlus Concanavalin A coated magnetic beads (Generon) were activated by washing and resuspension in Binding buffer (20mM HEPES pH 7.5, 10 mM KCl, 1 mM CaCl_2_, 1 mM MnCl_2_). 10 μL activated beads were added per sample and incubated at RT for 15 minutes. A magnet stand was used to isolate bead-cell complexes (henceforth ‘cells’), the supernatant was removed and cells resuspended in 50–100 μL Antibody buffer (20 mM HEPES pH 7.5; 150 mM NaCl; 0.5 mM Spermidine; 1× PIC; 0.05% Digitonin, 2 mM EDTA, 0.1 % BSA) and a 1:50 dilution of primary antibody (H3K27Ac (39034), Active Motif, H3K9me3 (C15410193) and H3K27me3 (C15410195), Diagenode; c-Jun (60A8) – 9165T, and JunD (D17G2) – 5000S, Cell Signalling; GATA3 sc-268 and TFAP2C sc-12762, Santa Cruz; TEAD4 CSB-PA618010LA01HU, Stratech and negative control rabbit IgG, sc-2027, Santa Cruz). Primary antibody incubation was performed on a nutator for 2 h at room temperature or overnight at 4 °C. Primary antibody was removed and secondary antibody (Guinea Pig anti-Rabbit IgG (Heavy & Light Chain) ABIN101961) diluted 1:50 in 50–100 μL Dig-Wash buffer, and added to the cells for 30 minutes incubation at RT. The supernatant was removed and cells were washed 3x in Dig-Wash buffer. Protein A-Tn5 transposase fusion protein (pA-Tn5) loaded with Illumina NEXTERA adapters (a kind gift from the Henikoff lab, via Madapura lab, diluted 1:250, or Epicypher CUTANA^®^ pAG-Tn5, diluted 1:10) was prepared in Dig-300 Buffer (0.05% Digitonin, 20 mM HEPES, pH 7.5, 300 mM NaCl, 0.5 mM Spermidine, 1× PIC) and 50–100 μL was added to the cells with gentle vortexing, followed by incubation at RT for 1 h. Cells were washed 3x in 800 μL Dig-300 buffer to remove unbound pA-Tn5. Next, cells were resuspended in 50–100 μL Tagmentation buffer (10 mM MgCl2 in Dig-300 Buffer) and incubated at 37 °C for 1 h. To stop tagmentation, 2.25 μL of 0.5 M EDTA, 2.75 μL of 10% SDS and 0.5 μL of 20 mg/mL Proteinase K was added to the sample and incubated overnight at 37 °C. To extract the DNA, 300 μL Phenol:Chloroform:Isoamyl Alcohol (PCI 25:24:1, v/v; Sigma-Aldrich) was added to the sample and vortexed. Samples were added to 5 PRIME™ Phase Lock Gel™ Light tubes and centrifuged for 3 minutes at 16,000 x g. Samples were washed in chloroform, centrifuged for 3 minutes at 16,000 x g and supernatant added to 100 % ethanol, chilled on ice and centrifuged 16,000 x g for 15 minutes. Pellets were washed in 100 % ethanol and allowed to air-dry, followed by resuspension in 30 μL 10 mM Tris-HCl pH8 1 mM EDTA containing 1/400 RNAse A. Libraries were indexed and amplified, using 21 μL DNA per sample and adding 25 μL NEBNext HiFi 2x PCR Master mix + 2 μL Universal i5 primer (10 μM) + 2 μL uniquely barcoded i7 primers (10 μM) in 0.2 ml PCR tube strips, using a different barcode for each sample. Cycling parameters were 72 °C for 5 minutes, 98 °C for 30 s, then 12 cycles of 98 °C for 10 s, 63 °C for 10 s, followed by 72°C for 1 minute followed by purification with 1X Agencourt AMPure XP beads as per the manufacturer’s instructions (Beckman Coulter).

### CUT&RUN (Cleavage Under Targets and Release Using Nuclease)

CUT&RUN was carried out as in ^31^. 500,000 hTSC were washed and resuspended in Wash buffer (20 mM HEPES pH 7.5, 150 mM NaCl, 0.5 mM Spermidine, plus PIC). Following Concanavalin A bead preparation (as for CUT&Tag), cells were incubated with 50 μL bead slurry for 15 minutes and supernatant removed. Cell-bead complexes (hereafter, ‘cells’) were incubated overnight with Antibody buffer (as for CUT&Tag) and primary antibodies (H3K27Ac (39034), Active Motif, H3K9me3 (C15410193) and H3K27me3 (C15410195), Diagenode; c-Jun (60A8) – 9165T, and JunD (D17G2) – 5000S, Cell Signalling; GATA3 sc-268 and TFAP2C sc-12762, Santa Cruz; TEAD4 CSB-PA618010LA01HU, Stratech and negative control rabbit IgG, sc-2027, Santa Cruz). Primary antibody was removed and cells washed 3x in Dig-Wash buffer and incubated with 1:200 pA-MNase (a gift from the Hurd lab) for 1 hr at 4 °C. Cells were washed 3x in Dig-wash buffer, followed by digestion (adding 2 μL 100mM CaCl2 to 100 μL sample) for 30 minutes at 0 °C, and the reaction stopped in STOP buffer (170 mM NaCl, 20 mM EGTA, 0.05% Digitonin, 100 ug/ml RNAse A, 50 ug/ml Glycogen) for 30 minutes at 37 °C. DNA was extracted using phenol/chloroform, as for CUT&Tag, and barcoded libraries constructed with the NEBNext^®^ Ultra™ II DNA Library Prep Kit for Illumina, using 12 cycles of PCR, followed by purification with 1X Agencourt AMPure XP beads.

### RNA Isolation and RT-qPCR

Total RNA was extracted with the AllPrep DNA/RNA Mini Kit (QIAGEN) and treated with TURBO DNA-free™ Kit (Ambion, AM1907) to remove contaminating genomic DNA. RNA (200ng) was retrotranscribed using Revertaid Reverse Transcriptase (Thermo Scientific, EP0441), and cDNA was diluted 1/50 for qPCRs using KAPA SYBR FAST (Sigma-Aldrich Co LTD, KK4610). RT-qPCR was carried out on a Roche LC480 for 40-45 cycles.

### RNA-seq

Prior to library construction, 10-100 ng total RNA was treated with the NEBNext^®^ rRNA Depletion Kit. Library construction was performed with the NEBNext^®^ Ultra II Directional RNA Library Prep Kit for Illumina^®^, according to the manufacturer’s protocol. RNA concentration and integrity was assessed using Bioanalyzer 2100 (Agilent Technologies, Santa Clara, CA), all hTSC samples had an RNA integrity number equivalent (RINe) value of >9.

### Guide RNA cloning into CRISPR–Cas9 plasmids

For CRISPR/Cas9 deletion of LTRs, sgRNA oligonucleotides (Integrated DNA Technologies) were designed to target upstream and downstream of the LTRs of interest, using Benchling (https://benchling.com) and annealed. Either the upstream or the downstream guide was cloned into plasmid LRG (Lenti_sgRNA_EFS_GFP; deposited by Christopher Vakoc (Addgene # 65656) which expresses GFP. The other guide was cloned into lentiCRISPR v2, deposited by Feng Zhang (Addgene plasmid # 52961). Clones were verified by Sanger sequencing (Source Bioscience). Guide sequences are listed in Supplementary Table S6.

### Lentivirus-mediated hTSC transduction and selection

Lentivirus was produced in 293T cells by quadruple transfection with CRISPR/Cas9 delivery vectors (see above) and the packaging plasmids psPAX2, (deposited by Didier Trono, Addgene plasmid # 12260) and pMD2.G (deposited by Didier Trono, Addgene plasmid # 12259) using FuGENE^®^ HD Transfection Reagent (Promega E2311). Viral supernatant was collected at 48 h and again at 72 h post transfection, pooled, filtered through 0.45 μm and either used fresh or aliquoted and stored at −80. hTSC cells were transduced with lentiviral supernatant supplemented with 4 μg/mL polybrene for 6 hours. Supernatant from lentivirus transfected with psPAX2, pMDG.2, empty LRG and empty lentiCRISPR v2 (i.e. no guide RNAs) was used alongside each new hTSC infection as a no-sgRNA control. 48 hours after transduction, GFP+ cells were sorted on a FACS Aria II and re-plated onto 6 or 12 well plates depending on cell number. 24-48 hours after sorting, cells were treated with Puromycin sulphate for 48 hours. DNA and RNA was extracted from resultant cell populations and genotyped for the presence of excisions. Successful excision pools were further analysed for the % of excised alleles and for the expression of genes nearby the targeted LTRs using qPCR and RT-qPCR. The number of independent RT-qPCR replicates used is visible on the respective plots – these were derived using different sgRNA sets or independent infections (as detailed in the figure legends and Supplementary Table S4), and/or from RNA collections at different passage numbers. Genotyping and RT-qPCR primers are listed in Supplementary Table S5.

### Endoglin ELISA

hTSC cell pools containing the LTR10A excision at ENG were seeded in parallel with no-sgRNA controls at 2.5 × 10^5^ cells per well of a six well plate and differentiated to SynT as described above. On day 6, media was collected and stored at −80. SynT cells were snap frozen for RT-qPCR analysis of differentiation markers and ENG expression. Media aliquots were frozen and subjected to ENG ELISA analysis with the Human Endoglin ELISA kit (Sigma-Aldrich RAB0171) according to the manufacturer’s instructions. Absorbance was measured at 450 nm.

### Transwell invasion assay

hTSC pools containing the LTR10A excision at ENG were seeded in parallel with no-sgRNA controls at 2.5 × 10^5^ cells per well of a collagen IV coated six well plate and differentiated to EVT as described above. On days 7-9 of differentiation, cells were treated with TrypLE and counted. A subset (~5%) of cells were tested for differentiation efficiency by HLAG staining and FACS to ascertain the proportion of HLAG+ cells, as described above. Invasion assays were carried out on either inserts coated with 75 μl Matrigel Basement Membrane Matrix (Fisher Scientific Uk Ltd, #11573620) diluted 1:1 with serum-free EVT medium; or commercial Matrigel pre-coated inserts (BioCoat Matrigel Invasion Chamber, VWR #734-1047), rehydrated by the addition of 500 μl serum-free D6 EVT medium in both upper and lower chambers for >1 hour at 37 °C/5% CO2; both utilising polycarbonate Transwell^®^ inserts (8.0 μm diameter pores, 6.5 mm diameter, Costar #CLS3464-48EA). Remaining cells (9,000 – 80,000, depending on the experiment) were resuspended in 500 μl serum-free D6 EVT medium and plated in the upper transwell insert chamber in duplicate. The lower chamber was filled with 500 μl media supplemented with 20% KSR. Cells were left to invade through the Matrigel for 48 or 72 h at 37 °C/5% CO2. Non-migrated cells were removed from the upper chamber using a cotton bud while cells that had migrated were fixed using 4% paraformaldehyde for 30 minutes and stained with 0.1% crystal violet solution for 15 minutes. Images were taken using a light microscope at 10x magnification.

### Primary sequencing data processing

High-throughput sequencing reads (2x 150 bp format; NovaSeq 6000) were trimmed using Trim_galore, with default settings. Mapping was done with either Bowtie2 ^88^ (for CUT&Tag, CUT&RUN and ChIP-seq; default settings) or Hisat2 ^89^ (for RNA-seq; with --no-softclip) to the reference genome of the species of origin: hg38, mm10 or rheMac10. Reads with MAPQ below 2 were discarded. Bigwig files were generated using deepTools ^90^ (bamCoverage tool with --binSize 200 --normalizeUsing CPM). Peak detection was performed using either MACS2 ^91^ (with -q 0.05 --broad) or SEACR ^92^ (with ‘norm’ and ‘relaxed’ options). RNA-seq gene raw counts or RPKM values were extracted using the RNA-seq pipeline in Seqmonk. DESeq2 ^93^ (default parameters) was used to perform differential expression analysis. SQuIRE ^94^ was used to measure ERV family-wide expression from RNA-seq data.

### Peak enrichment at repeat families

For a given ChIP-seq/CUT&Tag/CUT&RUN experiment, the number of peaks overlapping each Repeatmasker-annotated repeat family was compared with overlap frequencies across 1000 random controls (shuffled peaks, avoiding unmappable regions of the genome), yielding enrichment values and associated p-values. Significantly enriched repeat families had p<0.05, >2-fold enrichment, and at least 10 copies overlapped by peaks. Families were further selected by only keeping those for which >80% of ENCODE placental DNase-seq samples had an enrichment above 2-fold, and a median enrichment difference larger than 2 when compared to liver, lung and kidney ENCODE DNase-seq data.

### Transcription factor motif analysis

Motifs enriched at active TE families were identified using the AME tool of the MEME suite ^95^ (default parameters) and the 2020 JASPAR vertebrate database. Relevant motifs were selected based on the expression of the respective transcription factors in hTSC (>1 log2(FPKM)), followed by clustering to find redundant motifs and further selection based on literature searches. The FIMO tool was then used to extract motif locations and frequencies (default parameters).

### Human RNA-seq analysis

TE-derived promoters were identified by performing transcriptome assembly on primary cytotrophoblast data using Stringtie (with --rf and guided by the Gencode v38 annotation) ^96^, and intersecting the transcription start sites of multi-exonic transcripts with hTSC-active TE families. To evaluate putative enhancer effects, gene expression differences were determined based on the distance to the nearest H3K27ac-marked TE. JNK target genes were determined based on their distance to the nearest JUN binding peak. For all analyses, log2 fold differences were calculated using only genes that passed a minimal expression threshold (variable depending on dataset and normalisation strategy) in at least one of the two samples compared. Gene ontology analysis of differentially expressed genes after JNK inhibition was performed using topGO.

### Comparative analysis

Human TE orthologues were identified in non-human primates by performing a reciprocal liftOver. Human TSC RNA-seq data were merged to either mouse TSC or rhesus macaque TSC data based on one-to-one gene orthologues. For the human-mouse comparison, information about the proximity to hTSC-active TEs (from the selected families) and mTSC-active TEs (from the RLTR13D5 and RLTR13B families) was used to separate genes into different groups. For the human-macaque comparison, information about whether the nearest hTSC-active TE had a macaque orthologue was used.

### Statistical analyses

Wilcoxon tests with Benjamini-Hochberg correction for multiple comparisons, or ANOVA followed by Tukey post-hoc test were used, as specified in the figure legends.

## Data and code availability

CUT&Tag, CUT&RUN, ChIP-seq and RNA-seq data have been deposited in NCBI’s Gene Expression Omnibus under accession number GSE200763. Details of other datasets used can be found in Supplementary Table S7. All code associated with the manuscript is available in the GitHub repository https://github.com/MBrancoLab/Frost_2022_hTroph.

## Acknowledgements

We thank Jenna Kropp Schmidt and Tadeus Golos for sharing their macaque TSC RNA-seq data, Steve Henikoff, Pradeepa Madapura, Hong Wan and Paul Hurd for reagents, and Salvatore Papa for insights into the complexities of JNK activity. We also thank Gary Warnes for his gracious assistance with FACS.

