## Supplementary material for "Regulation of human trophoblast gene expression by endogenous retroviruses": Suplemmentary Figures

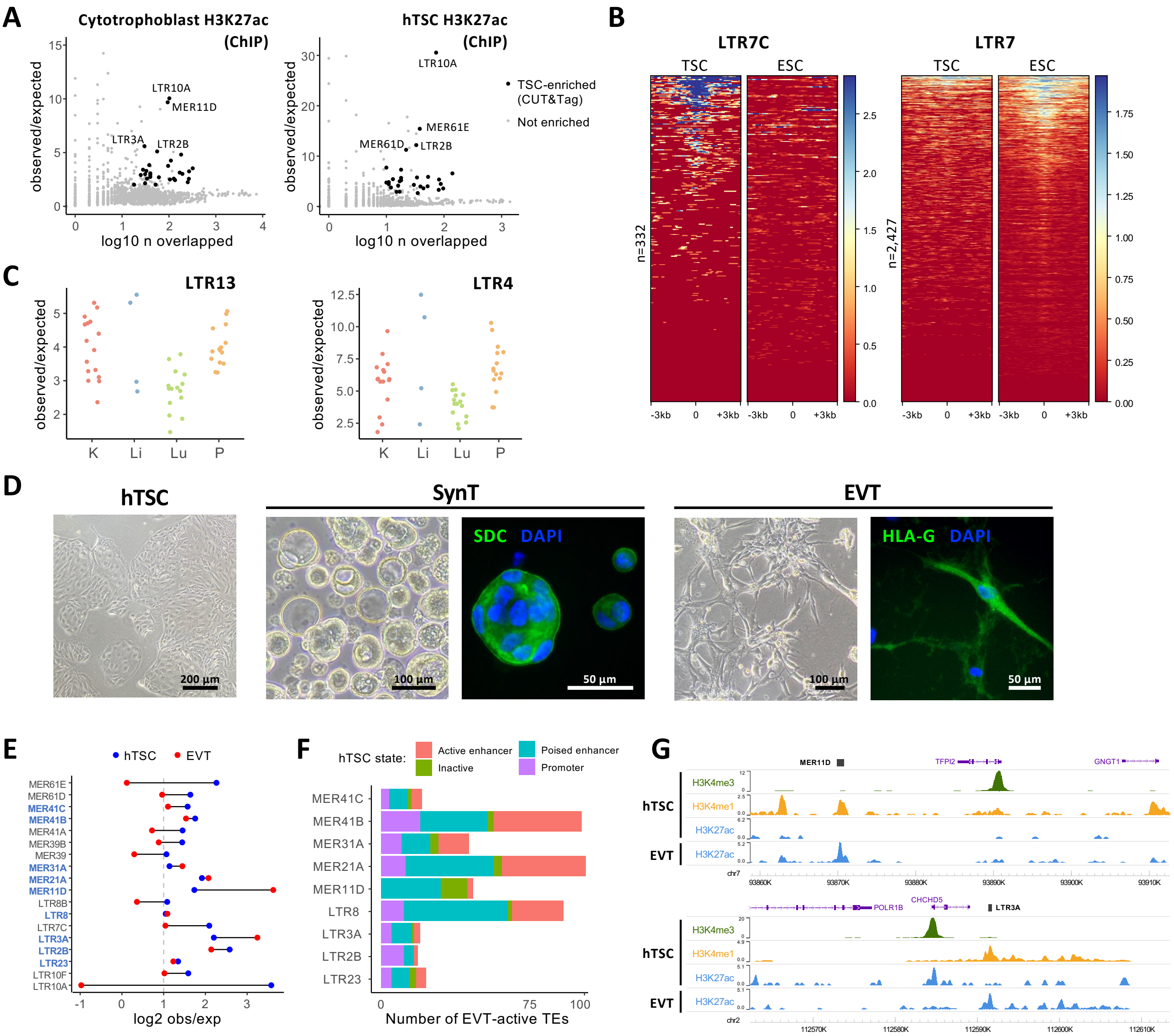


**Supplementary Figure 1 –** A) Enrichment for H3K27ac peaks for each repeat family in ChIP-seq data from primary cytotrophoblast or hTSCs. Families with significant enrichment in CUT&Tag data from hTSCs (Figure 1B) are highlighted. B) H3K27ac profiles of LTR7C and LTR7 families in hTSCs and hESCs. Each line represents an element in that family. C) Enrichment for DNase hypersensitive sites in LTR13 and LTR4 families, in kidney (K), liver (Li), lung (Lu) and placenta (P). Each datapoint represents a different ENCODE dataset. D) Representative phase contrast and immunofluorescence images for hTSCs and hTSC-derived SynT and EVT. Expression of the SynT marker SDC or EVT marker HLA-G is shown (green), with DAPI counterstain (blue). E) Enrichment for H3K27ac peaks in hTSC or EVT for all hTSC-enriched ERV families. Families that are also significantly enriched in EVT are highlighted in blue. F) Number of H3K27ac-marked ERVs in EVT, divided by their chromatin state in hTSCs: active enhancer (H3K4me1 + H3K27ac), poised enhancer (H3K4me1 alone), promoter (H3K4me3) or inactive (none of the three marks). G) Genome browser snapshots showing examples of ERVs that are in a poised state in hTSCs and become active in EVT.

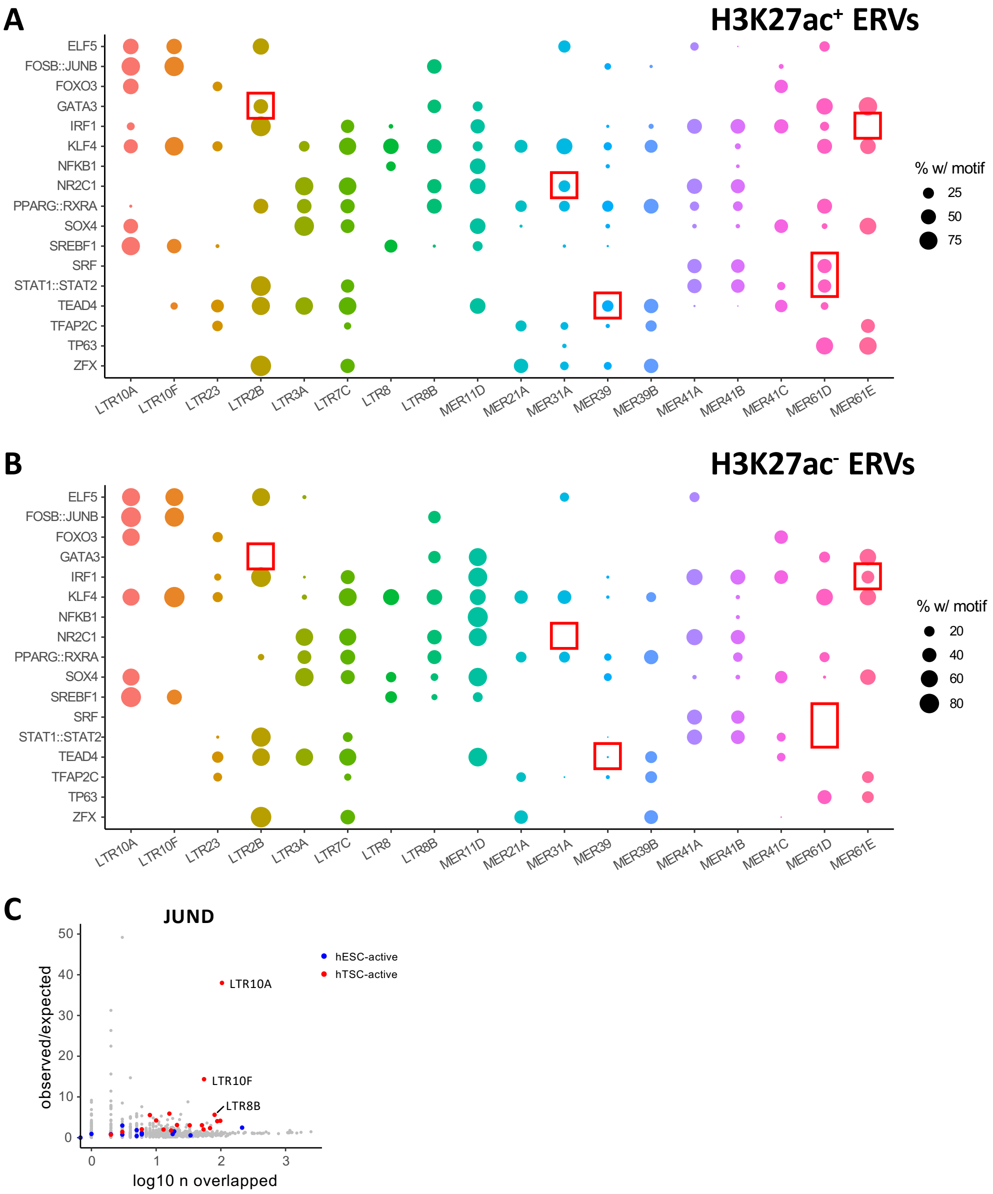


**Supplementary Figure 2 –** A) Proportion of H3K27ac-marked elements from all analysed ERV families bearing motifs for the transcription factors on the y axis. B) As in B, but for H3K27ac-negative elements. Particularly striking examples of differential motif enrichment between H3K27ac+ and H3K27ac- ERVs are highlighted. C) Repeat family-wide enrichment for peaks from JUND CUT&Tag data on hTSCs. H3K27ac-enriched families in hTSCs or hESCs are highlighted.

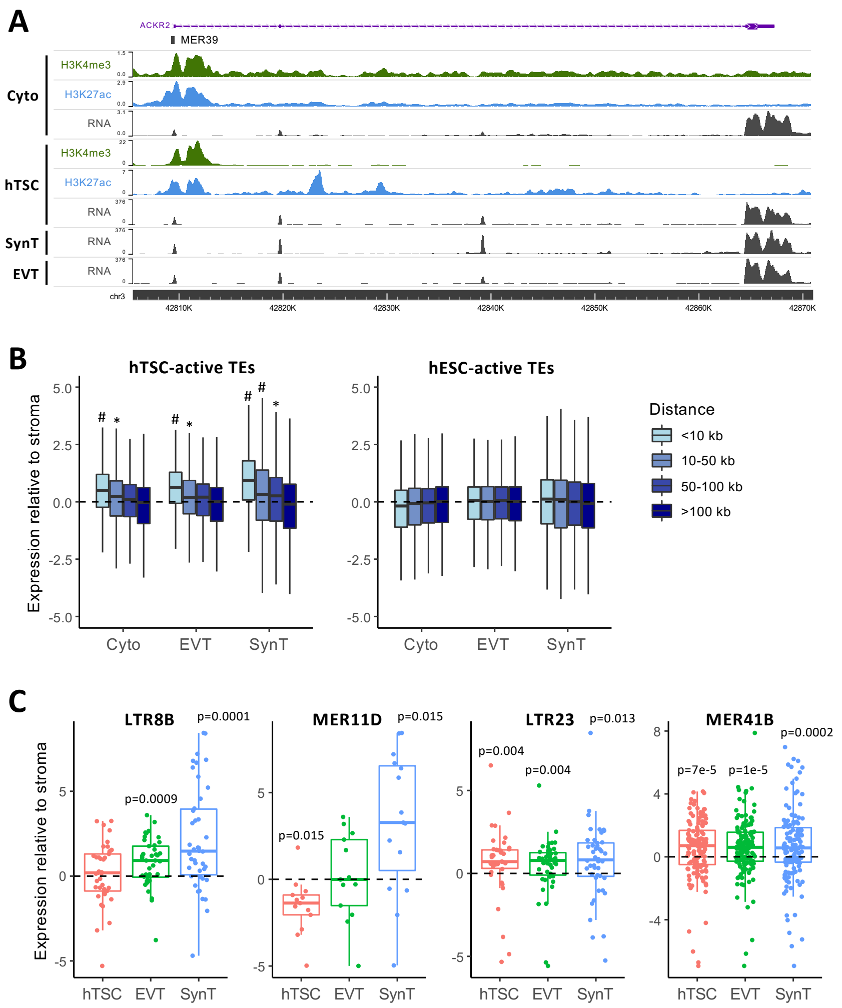


**Supplementary Figure 3 –** A) Genome browser snapshot showing an example of an ERV-derived gene promoter in primary cytotrophoblast. B) Gene expression in primary cytotrophoblast, EVT and SynT relative to placental stroma. Genes are grouped based on their distance to the nearest H3K27ac-marked TE in hTSCs. * $p<0.04$, # $p<3e-8$: difference to the '>100kb' group based on ANOVA and Tukey post-hoc test. C) Expression in hTSCs and hTSC-derived EVT and SynT relative to primary placental stroma for genes within 50kb of an H3K27ac-marked ERV of the indicated family. Genes are grouped based on their distance to the nearest H3K27ac-marked TE in hTSCs. P values are from Wilcoxon tests comparing each distribution to 0, with multiple comparisons correction.

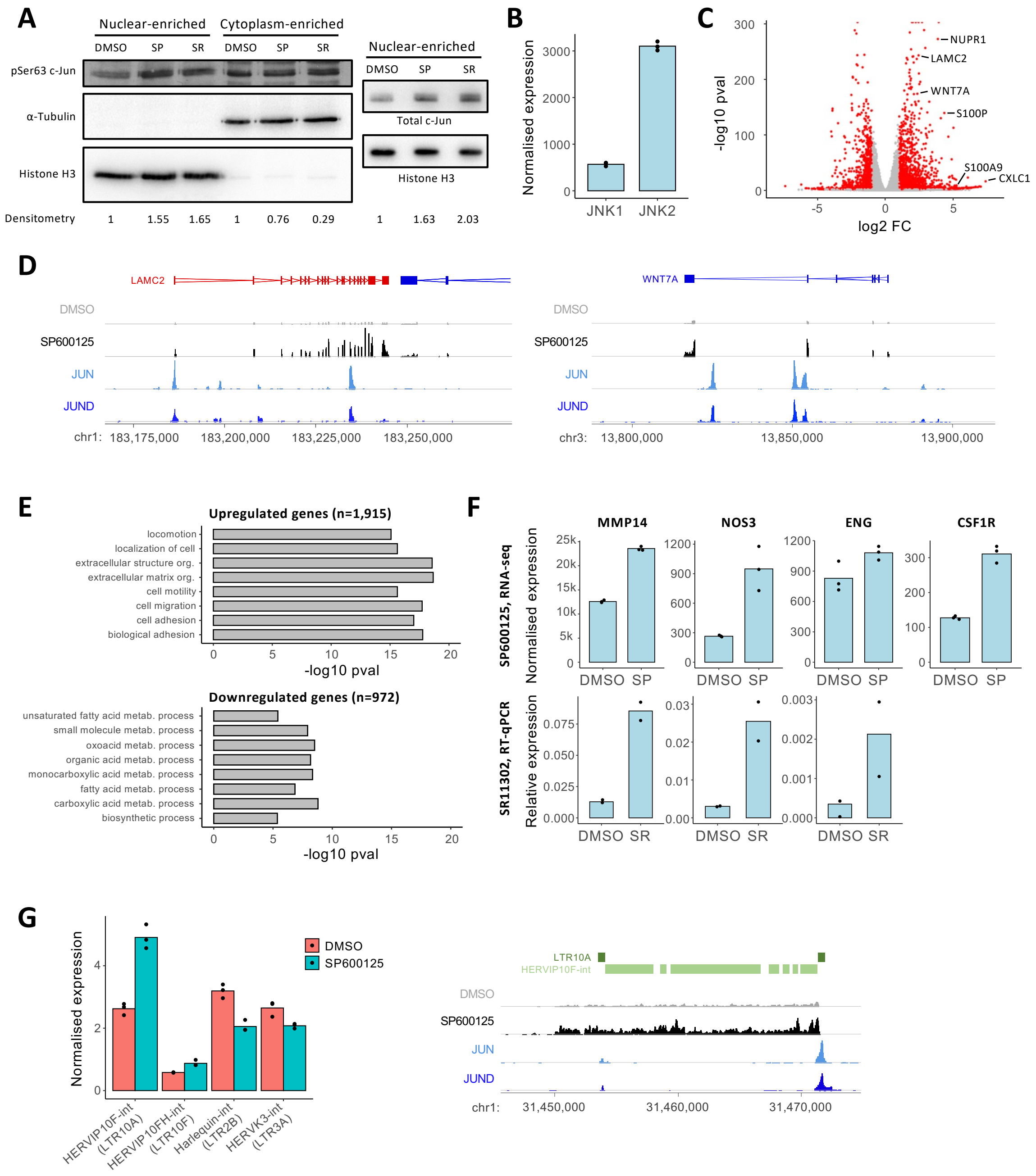


**Supplementary Figure 4 –** A) Western blot of phosphorylated c-Jun (pSer63 c-Jun) and total c-Jun in enriched nuclear or cytoplasmic fractions of cells treated with the AP-1 inhibitors SP600125 ('SP') or SR11302 ('SR'). Densitometry values normalized to the respective loading controls are shown below. Full blot images are in Supplementary Figure 7. B) Expression of *JNK1* (*MAPK8*) and *JNK2* (*MAPK9*) in hTSCs (data from RNA-seq). C) Volcano plot of RNA-seq data from cells treated with SP600125. Some genes implicated in cell migration are highlighted. D) Genome browser snapshots showing examples of genes highlighted in C, together with CUT&Tag data for JUN and JUND. E) Gene ontology biological processes enriched terms for genes up- or downregulated after treatment of hTSCs with SP600125. F) Expression of *MMP14* (a known AP-1 target), *NOS3*, *ENG* and *CSF1R* in hTSCs treated with SP600125 (data from RNA-seq) and/or SR11302 (data from RT-qPCR). G) Expression of internal ERV fragments driven by LTRs of H3K27ac-enriched families (in brackets) upon SP600125 treatment. An example of LTR10A-driven transcription of a proviral HERVIP10-F locus is shown on the right.

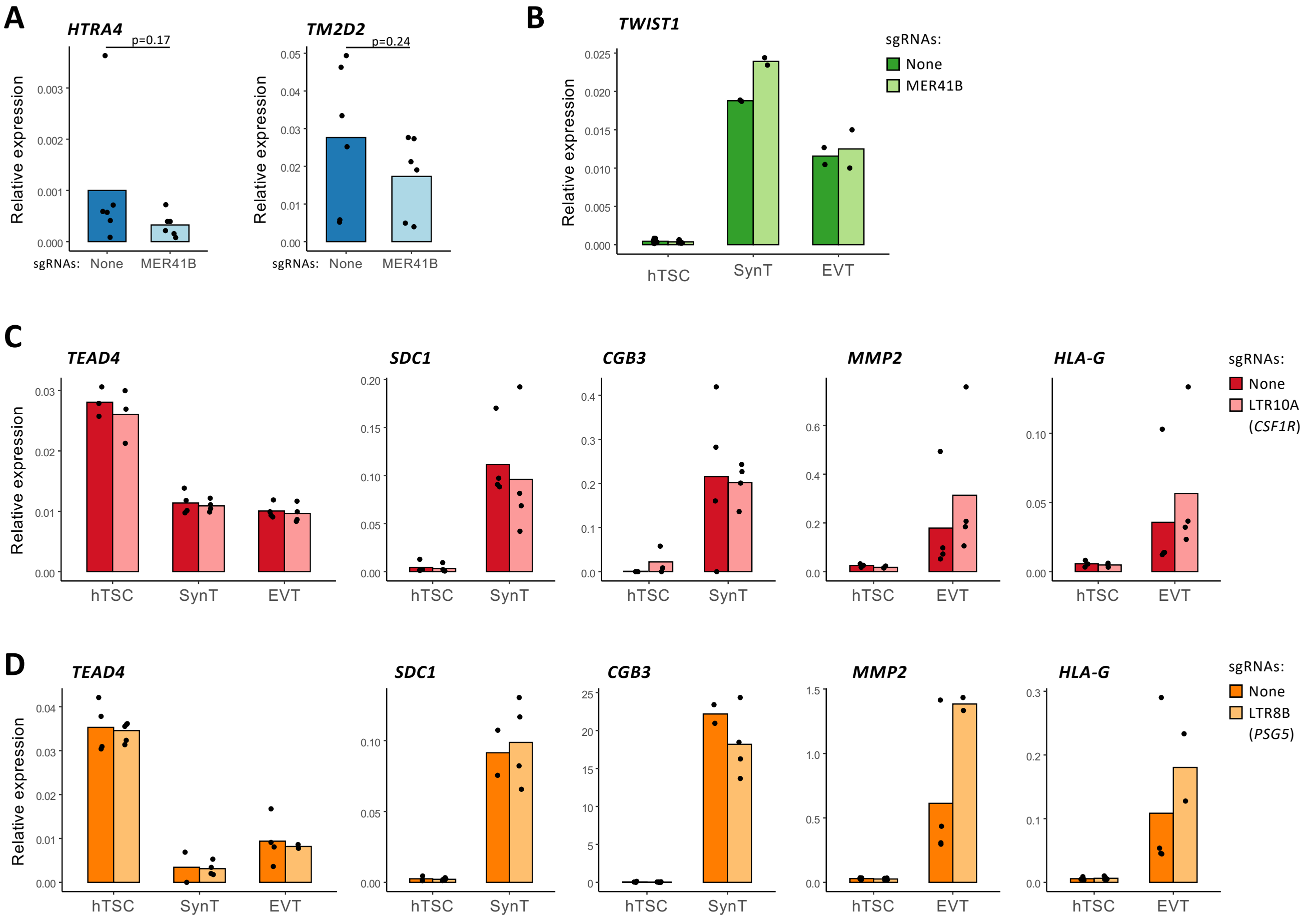


**Supplementary Figure 5 –** A) RT-qPCR data for *HTRA4* and *TM2D2* in hTSC populations treated with lentiviral CRISPR constructs carrying either no sgRNAs or sgRNAs that excise the MER41B element highlighted in Figure 5A. B) *TWIST1* expression in EVT and SynT derived from hTSC populations with no sgRNAs or sgRNAs that excise the MER41B element highlighted in Figure 5C. C) Expression of stem cell (*TEAD4*), SynT (*SDC1*, *CGB3*) and EVT (*MMP2*, *HLA-G*) markers in cell populations with no sgRNAs or sgRNAs that excise the LTR10A element highlighted in Figure 5B. D) As in C, but for the LTR8B element in Figure 5D. No significant differences were detected in any of the data after Wilcoxon tests with multiple comparisons correction.

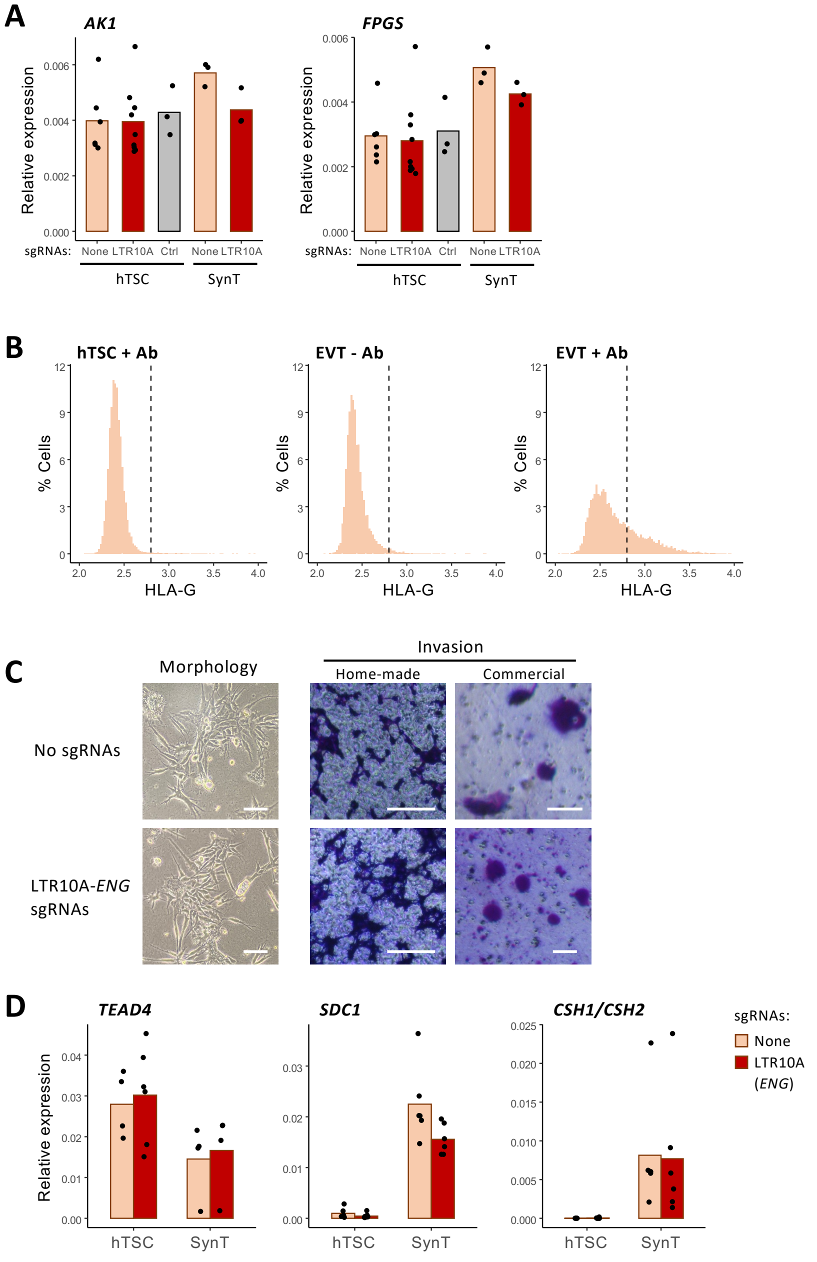


**Supplementary Figure 6 –** A) *AK1* and *FPGS* expression in hTSC or hTSC-derived SynT populations carrying lentiviral CRISPR constructs with no sgRNAs, sgRNAs that excise the LTR10A element highlighted in Figure 6A, or sgRNAs that excise a control region also highlighted in Figure 6A. B) Representative FACS profiles for HLA-G immunolabelling in hTSCs and hTSC-derived EVTs. The middle panel is a no-antibody control. C) Images of cells in culture (to assess morphology) or crystal violet-stained cells after an invasion assay using Matrigel-coated chambers (either home-made chambers or from a commercial provider). Scale bars are 100 µm. D) Expression of stem cell (*TEAD4*) and SynT (*SDC1*, *CSH1/CSH2)* in cell populations with no sgRNAs or sgRNAs that excise the LTR10A-*ENG* element. No significant differences were detected in any of the qRT-PCR data after Wilcoxon tests with multiple comparisons correction.

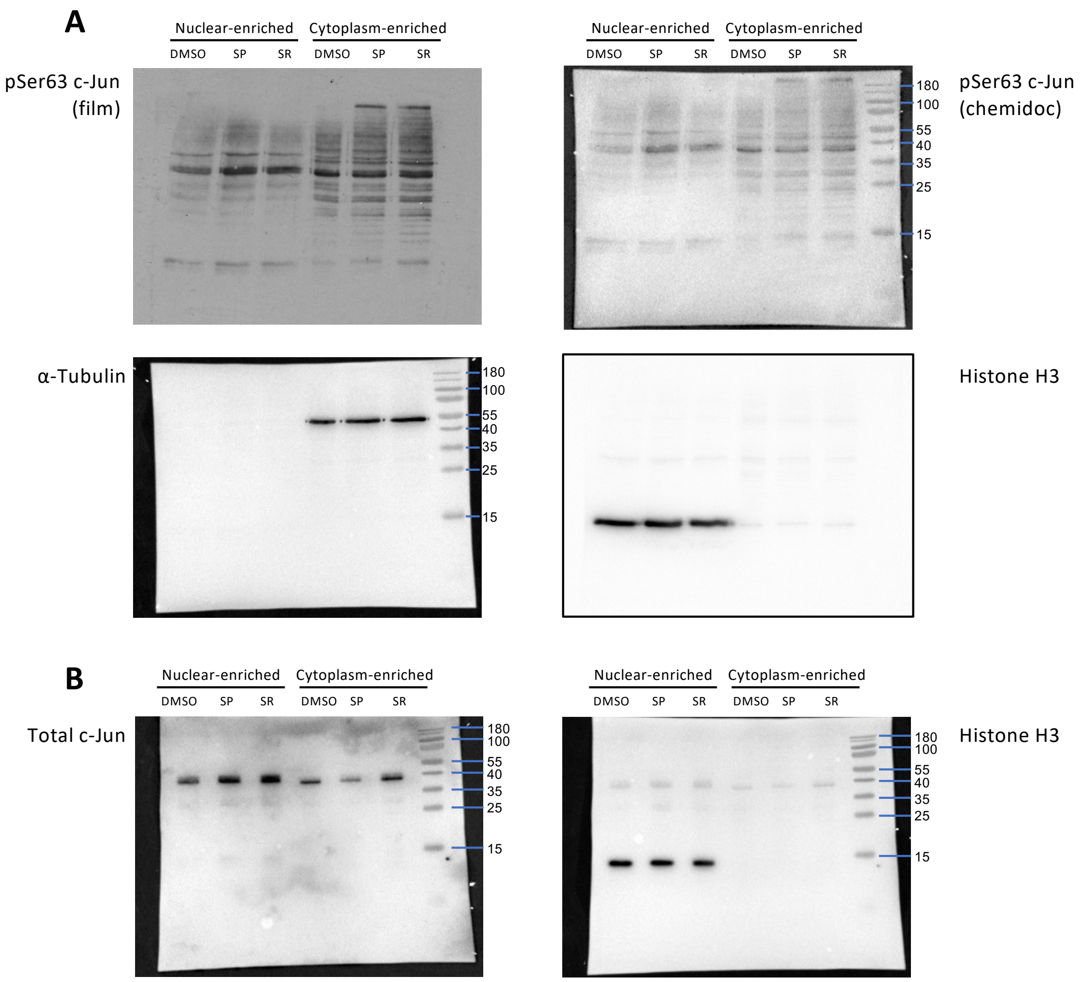


**Supplementary Figure 7 –** Uncropped images of the western blot data shown in Supplementary Figure 4A. Molecular weight (kDa) markers are shown (PageRuler™ Prestained Protein Ladder, 10 to 180 kDa, catalog number: 26616, Thermofisher). A) Images from blot for phosphorylated c-Jun. B) Images from blot for total c-Jun. Note that the c-Jun bands are also visible on the histone H3 image, as antibodies were incubated sequentially.
